# Transferrin-dependent crosstalk between the intestinal tract and commensal microbes contributes for immune tolerance

**DOI:** 10.1101/2020.03.02.972281

**Authors:** Xiaopeng Tang, Mingqian Fang, Kuanhong Xu, Ruomei Cheng, Gan Wang, Zhiyi Liao, Zhiye Zhang, James Mwangi, Qiumin Lu, Ren Lai

## Abstract

Crosstalks between gastrointestinal tract and commensal microbes regulate immune tolerance and maintain host intestinal homeostasis. However, molecular events that regulate the crosstalks remain poorly understood. Here, we show that microbial products (lipopolysaccharide, lipoteichoic acid and DNA) up-regulate host transferrin, an iron supplier of commensal bacteria, to induce host’s immune tolerance by negatively regulating toll-like receptor (TLR) signaling. Transferrin level in germ-free and broad-spectrum antibiotics-treated mice is much less than that in normal mice. Transferrin is found to silence TLR signaling complex by directly interacting with CD14, a co-receptor of many TLRs. Transferrin knock-down impaired host tolerogenic responses as well as broad-spectrum antibiotics treatment. Our findings reveal that commensal bacteria up-regulate and beneficially use host transferrin as a negative regulator of TLR signaling to shape host immunity and contribute for intestinal tolerance.

## Introduction

The animal gastrointestinal tract is colonized by various microorganisms, including bacteria, fungi, and viruses(Frosali et al., 2015; Hooper and Gordon, 2001), with commensal bacteria forming the largest population of intestinal microbiota. The animal host and its intestinal microbial flora impact one another and function together as a complex ecological system(Rolfe, 1984). On the one hand, healthy intestines have the ability to shape microbiota and limit colonization of the intestinal tract by harmful bacteria through symbiotic relationships(Hooper and Macpherson, 2010; MacPherson and Uhr, 2004). Host colon mucus layers create a physical barrier composed of mucin glycoproteins that affect host-microbial interactions by separating bacterial flora and intestinal epithelial cells(Johansson and Hansson, 2011; Johansson et al., 2011). Moreover, hosts secrete antibacterial factors such as defensins, C-type lectins, lysozyme, phospholipase A2, and IgAs to regulate microbial growth(Benckert et al., 2011; Gallo and Hooper, 2012). On the other hand, intestinal microbiota play crucial roles in shaping the immune functions of their hosts(Mowat, 2018). Intestinal microbiota can stimulate the development of both local and systemic host immunity(Tlaskalova-Hogenova et al., 2011). Significantly down-regulated immune responses and substantially smaller lymphoid organs have been found in germ-free animals(Carter and Pollard, 1971; Tlaskalova et al., 1970), whereas microbial colonization can increase systemic immunological capacity by elevating immunoglobulin and antibody levels and changing mucosal-associated lymphocyte tissues and cell populations(Stepankova et al., 1998; Tlaskalovahogenova et al., 1981; Tlaskalovahogenova et al., 1983). One main characteristic of the intestine is its ability to maintain tolerance to microbial antigens, thus showing a symbiotic host relationship(Guarner and Malagelada, 2003; Mowat, 2018). Long evolution by mutually adapting and selecting each other creates the close symbiosis of the microbiota and its hosts. The symbiotic relationship maintains a constant homeostasis by perfectly regulating the microbial load and the immune response generated against it in the healthy human intestine, while dysbiosis of intestinal flora may result in various pathological conditions(Abraham and Cho, 2009; Sartor, 2008; Xavier and Podolsky, 2007).

Hosts can identify invading micro-organisms and trigger the activation of innate immunity, leading to the development of antigen-specific adaptive immunity via recognition of microbial components by host toll-like receptors (TLRs)(Takeda and Akira, 2001). However, commensal bacteria use a similar mechanism to enhance colonization of the gut and establish host-microbial tolerance(Frosali et al., 2015). For example, certain bacteria, such as *Bacteroides fragilis*, establish host-microbial symbiosis via polysaccharide A, which activates the TLR2 pathway and induces the production of anti-inflammatory cytokine IL-10 in T-cells to restrain Th17 responses(Round et al., 2011). Tolerance to cell-wall components of gram-negative and gram-positive bacteria (e.g., lipopolysaccharide (LPS) and lipoteichoic acid (LTA)) has also been reported(Astiz et al., 1995; Seeley and Ghosh, 2017; Yoon et al., 2017). Given that human gut microbiome comprises ten trillion diverse symbionts (>50 bacterial phyla and >1000 bacterial species) representing one of the most densely populated ecosystems known, immune silencing might be a microbe-intrinsic feature and some products derived from microbiome may have the ability to facilitate host tolerance. However, it is unknown that the mechanism to silence host immunity by microbiome-derived products and if the products have common immunosuppressive pathway. Although commensal gut bacteria communities vary considerably with time and environmental parameters(Engel and Moran, 2013; Wong et al., 2013) and may lead to variable gut microbiota-host interplay, the host generally maintains gut immune homeostasis. How does the inconstant interplay consequently achieve the general immune homeostasis or tolerance? Here we reported that microbiome-derived products (LPS, LTA and DNA) induce the host’s immune tolerance via up-regulating host transferrin to silence TLR Signaling.

## Results

### Transferrin is a potential contributory factor for intestinal tolerance

Mass spectrometry-based label-free quantification was used to analyze differential protein expression in the plasma of specific pathogen-free (SPF) and germ-free (GF) mice. As illustrated in Figure S1, transferrin showed decreased protein expression in GF mice. Based on enzyme linked immunosorbent assay (ELISA), the average transferrin concentration in the plasma of GF mice (n = 10, male 5; female 5) was 1.84 mg/ml (SD 0.32), whereas that in SPF mice (n = 10, male 5; female 5) was 2.237 mg/ml (SD 0.27) (Figure 1A). Plasma transferrin levels returned to normal when the GF mice were fed in the SPF environment for four weeks (Figure 1A). Decreased levels of transferrin were also observed in the liver and spleen of GF mice (Figures 1B and 1C). As illustrated in Figure 1D, transferrin levels in different segments of gut tissue (e.g., duodenum, jejunum, ileum, and colon (D, J, I, and C2)) of GF mice were lower than levels in SPF mice. This was consistent with the lower levels of aldehyde dehydrogenase 1 family member A2 (ALDH1A2), CCL22, TGF-β, and IL-10 observed in GF mice, which are known to play key roles in maintaining intestinal tolerance(Esterhazy et al., 2019). After treatment with broad-spectrum antibiotics (Abs) to reduce gut microbiota, decreases in transferrin and corresponding ALDH1A2, CCL22, TGF-β, and IL-10 levels were also observed in the gut of Abs-treated mice (Figure 1E).

**Figure 1.**
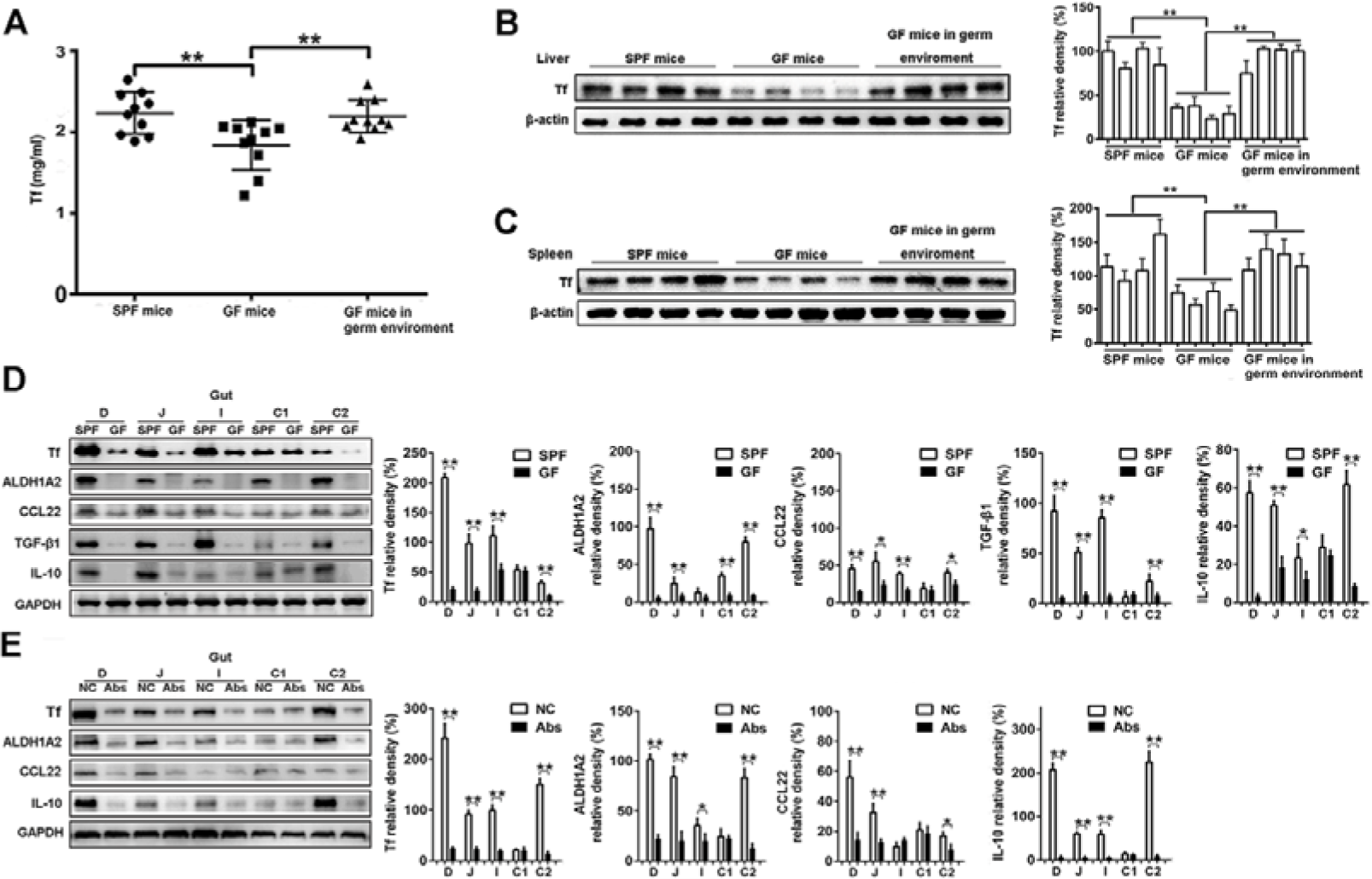
Transferrin is a potential contributory factor for intestinal tolerance dependent on gut microbiota. **(A)** Transferrin concentration in plasma of germ-free (GF) and specific pathogen-free (SPF) (control) mice determined by ELISA. Data represent means ± SD (n = 10), \**p* < 0.05, \*\**p* < 0.01 by one-way ANOVA with Dunnett’s *post-hoc* test. Western blotting (left) and quantification (right) analyses of transferrin levels in protein extracts from liver **(B)** and spleen **(C)** of SPF and GF mice. β-actin was used as the control. Data represent means ± SD (n = 10), \**p* < 0.05, \*\**p* < 0.01 by one-way ANOVA with Dunnett’s *post-hoc* test **(B, C). (D-E)** Transferrin, ALDH1A2, CCL22, TGF-β, and IL-10 levels in different gut tissue segments (duodenum, jejunum, ileum, caecum, and colon (D, J, I, C1, and C2)) of SPF (NC), GF (**D**), and broad-spectrum antibiotic (Abs)-treated SPF (**E**) mice were determined by Western blotting (left) and quantification (right) analyses. GAPDH was used as the control. Data represent means ± SD (n = 8), \*\**p* < 0.01 by unpaired *t-*test **(D, E)**.

We next tested the effects of microbial products (i.e., LPS, LTA, and DNA) on the expression of transferrin in human monocytic cell line (THP-1), human umbilical vein endothelial cells (HUVECs), mouse normal embryonic liver cell line (BNL CL.2), and mouse splenocytes. Western blot analysis (Figures 2A-D and Figures S2A-D), quantitative real-time polymerase chain reaction (qRT-PCR) (Figures S5A-D), and ELISA (Figures S5E-G) indicated that transferrin was up-regulated by LPS in a dose-dependent manner in all cells. Similarly, LTA (Figures 2E-G, Figures S3A-D, Figures S6A-D, and Figures S6E-H) also up-regulated transferrin in these cells. In addition, bacterial DNA from *Escherichia coli, Staphylococcus aureus*, and *Listeria monocytogenes*, but not mouse DNA, up-regulated transferrin in these cells (Figure 2H, Figure S4, Figures S7A-D, and Figures S7E-H). Importantly, Western blot (Figure 2I and Figure S8), qRT-PCR (Figure S9), and ELISA (Figure S10) analyses indicated that all transferrin up-regulation induced by LPS, LTA, and bacterial DNA was inhibited by the NF-κB inhibitor pyrrolidine dithiocarbamate (PDTC).

**Figure 2.**
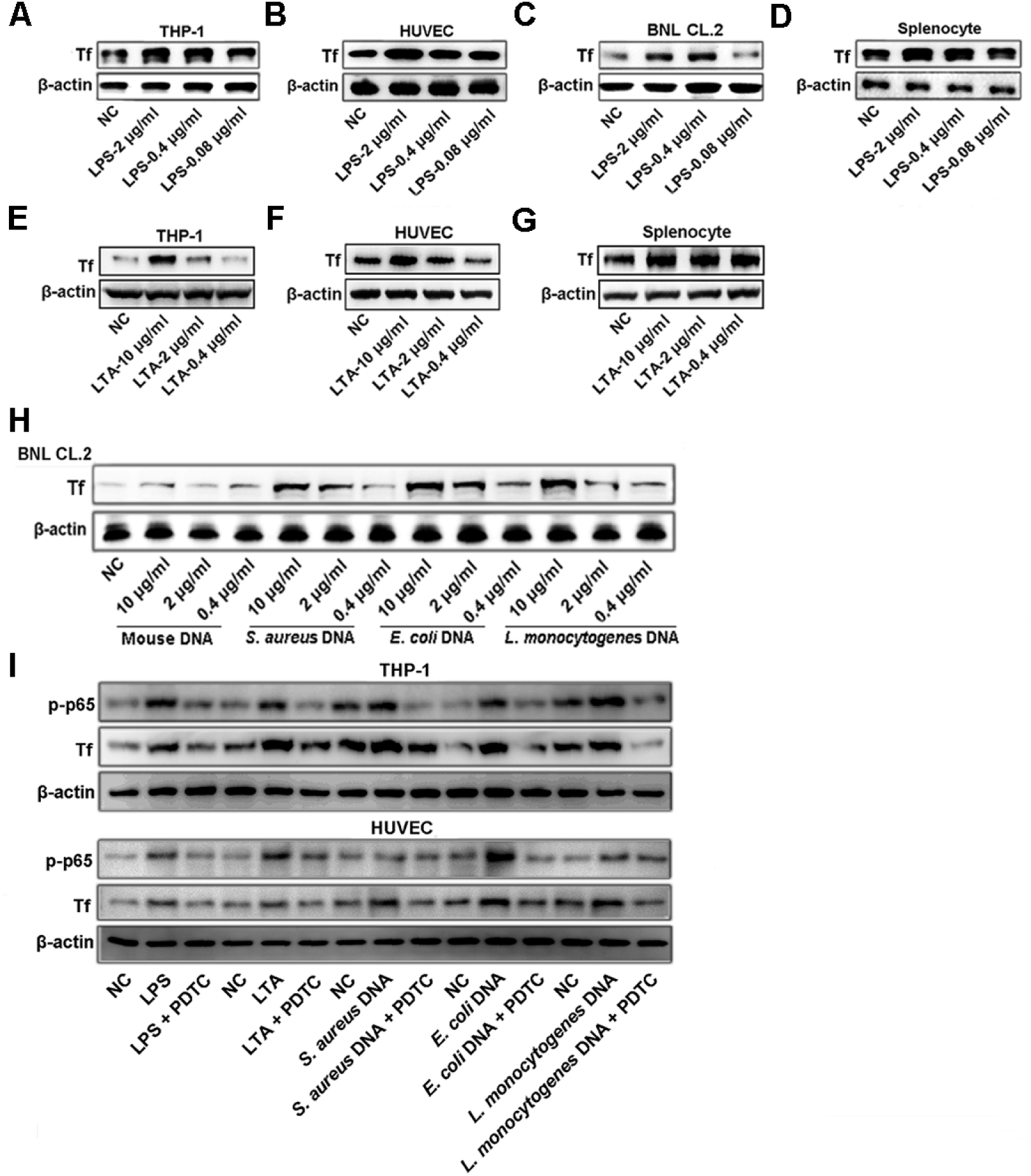
Microbial products up-regulate host transferrin. Effects of LPS **(A-D)**, LTA **(E-G)**, and bacterial DNA **(H)** on transferrin expression in THP-1 cells, HUVECs, BNL CL.2 cells, or splenocytes were analyzed by Western blotting. THP-1 cells and HUVECs were treated with LPS (2 μg/ml), LTA (10 μg/ml), *E. coli* DNA (10 μg/ml), *S. aureus* DNA (10 μg/ml), or *L. monocytogenes* DNA (10 μg/ml) after pyrrolidine dithiocarbamate (PDTC) pretreatment (20 μM) for 30 min. Transferrin expression was analyzed by Western blotting **(I)**. β-actin was used as the control. Tf: transferrin.

### Cluster of differentiation 14 (CD14) is a direct target of transferrin

Surface plasmon resonance (SPR) analysis revealed that transferrin directly interacted with CD14 (Figure 3A). The association (*Ka*), dissociation (*Kd*), and equilibrium dissociation constants (*KD*) of the interaction between transferrin and CD14 were 2.4 × 10^4^ M^−1^s^−1^, 3.3 × 10^−4^ s^−1^, and 14 nM, respectively. Moreover, native gel shift assay showed complex formation between transferrin and CD14 (Figure 3B). Confocal microscopy showed the colocalization of CD14 and FITC-labeled transferrin on the cell membrane of THP-1 cells, whereas transferrin antibody blocked the colocalization (Figure 3C).

**Figure 3.**
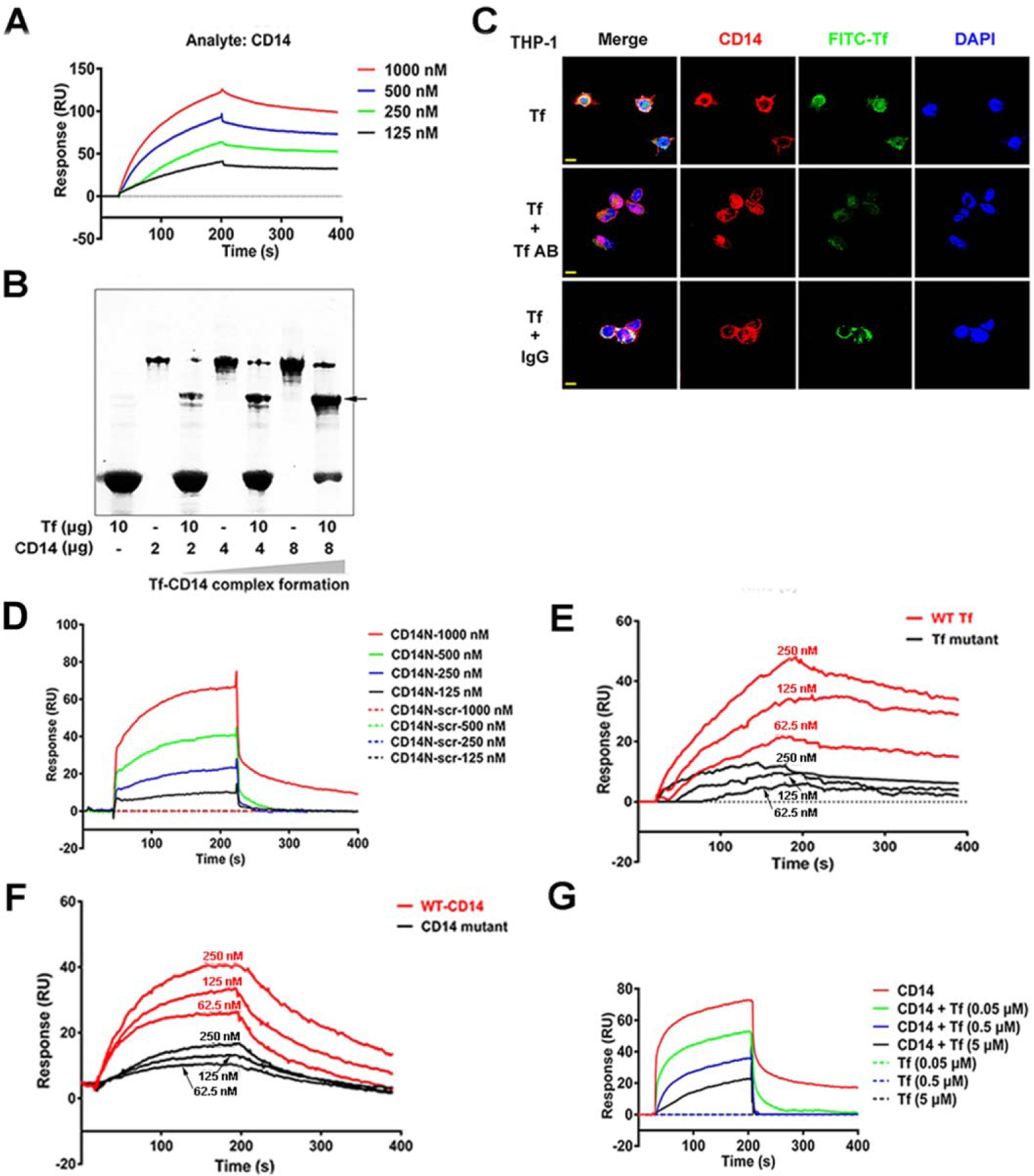
Interaction between transferrin and CD14. **(A)** SPR analysis of interaction between transferrin and CD14. **(B)** Native gel shift analysis of interaction between transferrin (10 μg) and CD14 (2, 4, and 8 μg). Colocalization of CD14 and FITC-labeled transferrin on cell membranes of THP-1 cells **(C)** was analyzed by confocal microscopy. Cell nuclei were labeled by DAPI. Scale bar represents 10 μm. Images are representative of at least three independent experiments. **(D)** SPR analysis of interaction between transferrin and LPS binding region of CD14 N-terminus (CD14N) and CD14N-scr (scrambled control of CD14N). **(E)** SPR analysis of interaction between wild-type transferrin (WT-Tf) or transferrin mutant (R663A and K664A, Tf mutant) and CD14. **(F)** SPR analysis of interaction between transferrin and wild-type CD14 (WT-CD14) or CD14 mutant (D44A, S46A and Q50A). **(G)** Effects of transferrin on interaction between LPS and CD14 by SPR analysis. Tf: transferrin.

The docking model of the transferrin-CD14 complex (Figure S11) suggested that the LPS binding region (CD14N: ELDDEDFRCVCNFSEPQPDWSEAFQCVSAVEVEIHAGGLN) located at the N-terminus of CD14 was responsible for the transferrin-CD14 interaction, as confirmed by SPR analysis (Figure 3D). We identified two key residues of transferrin (R663 and K664) that likely play key roles in the transferrin-CD14 interaction. The mutant of transferrin (R663A and K664A) was thus constructed (Figures S12A, B, and E). Notably, the mutant exhibited weak interaction with CD14 (Figure 3E). Furthermore, the docking model also suggested that three key residues of the LPS-binding region of the CD14 N-terminus (D44, S46, and Q50) may participate in the transferrin-CD14 interaction. The corresponding mutants (D44A, S46A, and Q50A) of CD14 (Figures S12C, D and E) exhibited weak interaction with transferrin compared with wild-type CD14 (Figure 3F). Importantly, the interaction between CD14 and LPS was blocked by transferrin (Figure 3G).

### Transferrin inhibits TLR4 activation induced by LPS

As illustrated in Figures 4, transferrin blocked LPS-induced phosphorylation of transforming growth factor-β (TGF-β)-activating kinase 1 (TAK1), IκB (inhibitory subunit of nuclear factor κB (NF-κB)) kinase α (IKKα), IκBα, and NF-κB p65 of the myD88-dependent pathway and TRAF-associated NF-κB activator (TANK)-binding kinase 1 (TBK1) and interferon regulatory factor 3 (IRF3) of the myD88-independent pathway in THP-1 cells.

**Figure 4.**
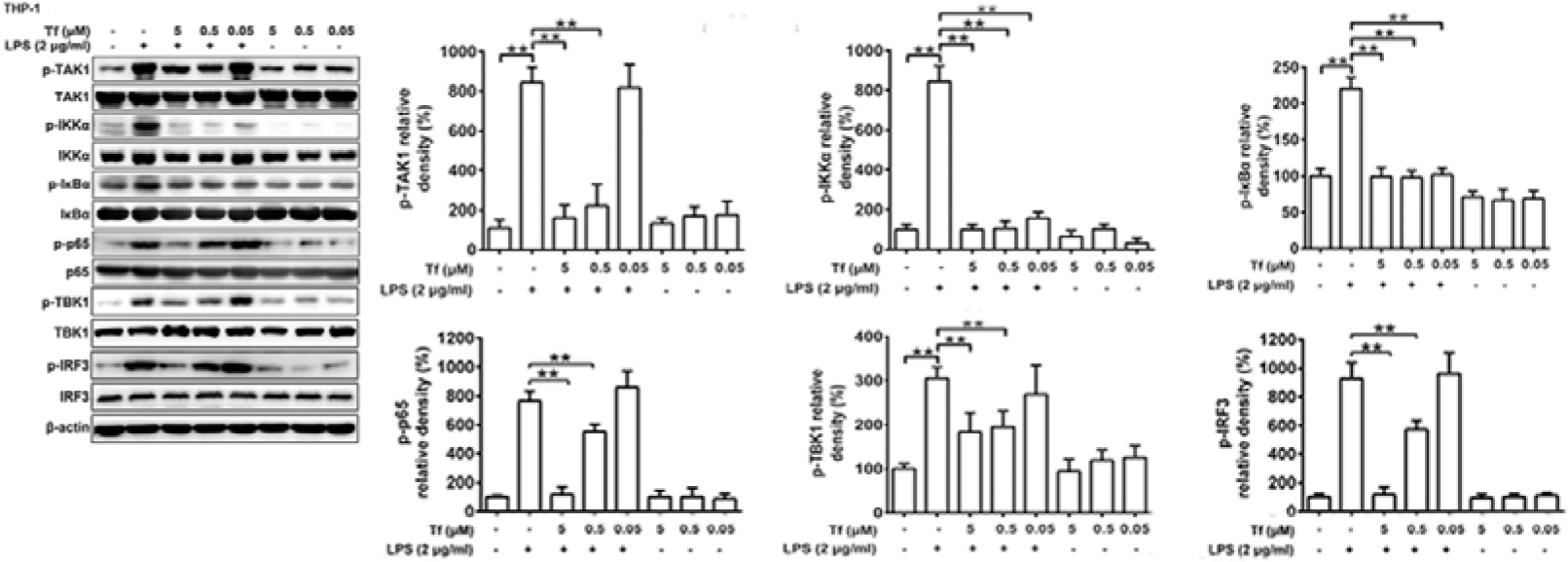
Transferrin inhibits TLR4 activation. THP-1 cells were stimulated in presence or absence of transferrin by LPS for 8 h. Total and phosphorylated TAK1, IKKα, IκBα, and p65 of the myD88-dependent pathway and TBK1 and IRF3 of the myD88-independent pathway in THP-1 cells were analyzed by Western blotting, respectively. Corresponding quantifications are shown on right (six panels). β-actin was used as loading control. Data represent means ± SD of five independent experiments, \**p* < 0.05, \*\**p* < 0.01 by one-way ANOVA with Dunnett’s *post-hoc* test. Tf: transferrin.

TLR4 stimulation by LPS induces the release of many cytokines to activate potent immune responses(Lu et al., 2008). As transferrin inhibits TLR4 activation evoked by LPS, as shown above, we investigated transferrin’s effects on cytokine release stimulated by LPS. As an iron carrier, transferrin exists in plasma in both the ferric iron-bound (holo-transferrin) and unbound states (apo-transferrin). As illustrated in Figures S13-17, both apo- and holo-transferrin showed similar inhibition of the LPS-induced production of TNF-α, IL-6, IFN-β, or TGF-β in human peripheral blood mononuclear cells (PBMCs), polymorphonuclear neutrophils (PMNs), mice bone marrow dendritic cells (BMDCs), THP-1 cells, and HUVECs in a dose-dependent manner. Importantly, the blockage of transferrin-transferrin receptor (TfR) interaction using anti-TfR antibody (TfR AB) had no effect on the inhibition of cytokine release elicited by transferrin, suggesting that the inhibition is independent of TfR (Figures S13-17).

### Transferrin overexpression and knockdown attenuates and aggravates inflammatory response induced by LPS, respectively

Transferrin overexpression (PLP-Tf) and knockdown mice (RNR-Tf) were used to further elucidate the role of transferrin in inflammatory response (Figure S18). As illustrated in Figures 5A-D, the plasma levels of inflammatory factors induced by LPS, including TNF-α, IL-6, IL-1β, and IFN-β, decreased following transferrin overexpression and in mice directly administered with an intravenous injection of exogenous transferrin, but were exacerbated by transferrin knockdown. Effects of transferrin on inflammatory injury induced by LPS were evaluated by histological examination (Figures 5E and F). Transferrin overexpression and exogenous transferrin administration alleviated LPS-induced liver injury, whereas the injury was aggravated by transferrin knockdown (Figure 5E). Apoptosis detection indicated that transferrin overexpression and exogenous transferrin administration inhibited LPS-induced apoptosis, whereas transferrin knockdown promoted LPS-induced apoptosis (Figure 5F).

**Figure 5.**
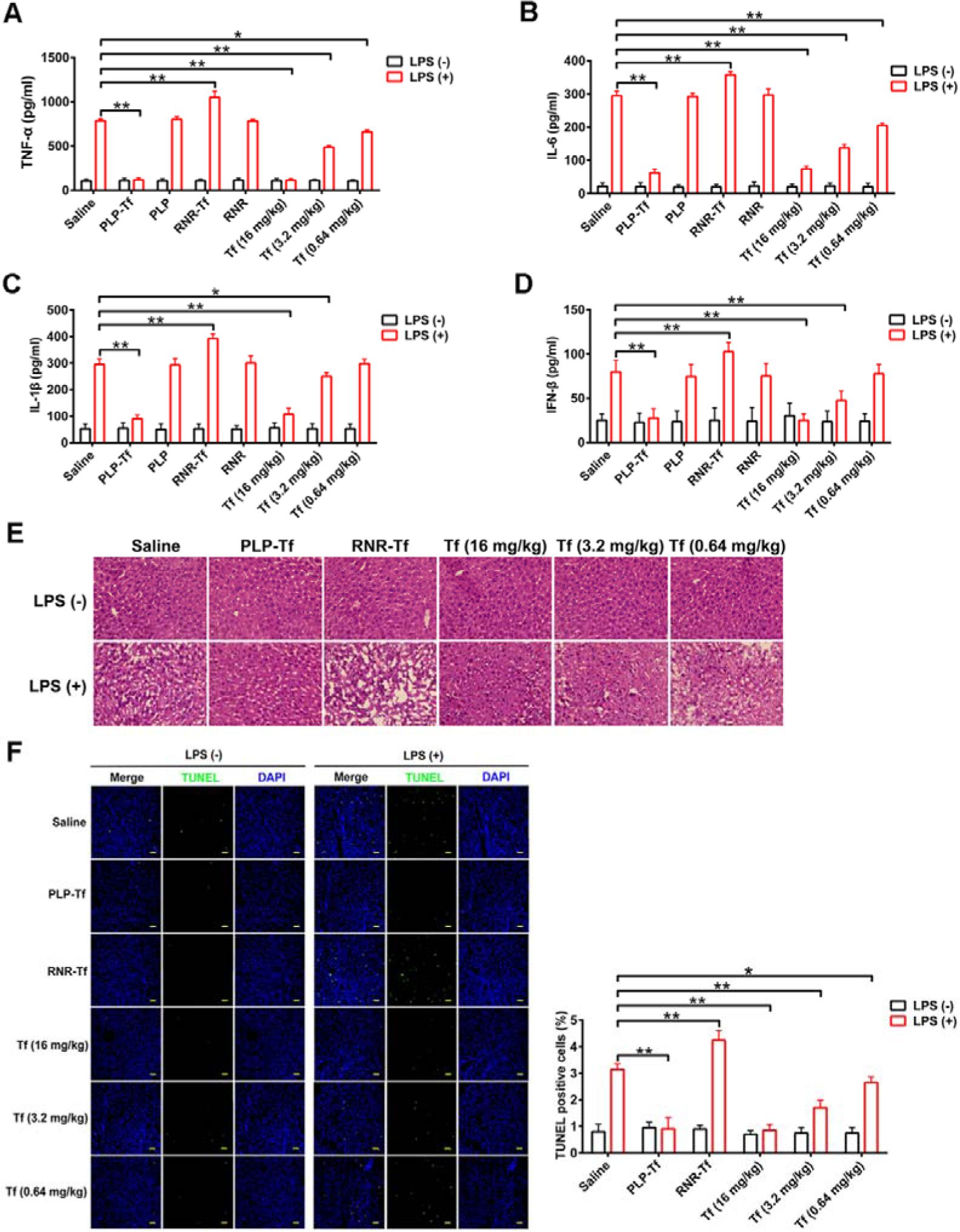
Effects of transferrin on inflammatory response induced by LPS *in vivo.* LPS (750 μg/kg) was injected into the tail vein of mice, including transferrin overexpression (PLP-Tf), knockdown (RNR-Tf), or their blank (PLP and RNR), to induce an inflammatory response for 2 h. In transferrin-treated group, LPS injection was performed after transferrin administration through the tail vein for 20 min. Plasma TNF-α **(A)**, IL-6 **(B)**, IL-1β **(C)**, and IFN-β **(D)** levels were determined by ELISA. **(E)** LPS-induced liver injuries in all mouse groups were determined by hematoxylin and eosin (H&E) staining. **(F)** Apoptosis induced by LPS was also evaluated using an apoptosis detection kit and the corresponding quantifications are shown on right. Data represent means ± SD (n = 10), \**p* < 0.05, \*\**p* < 0.01 by one-way ANOVA with Dunnett’s *post-hoc* test **(A-F)**. Tf: transferrin.

### Transferrin down-regulation impairs host tolerogenic responses

CD103^+^CD11b^+^ dendritic cells (DCs), regulatory T cells (Tregs), and regulatory B cells (Bregs) have been implicated in gut tolerogenic responses(Esterhazy et al., 2019; Mortha et al., 2014). Here, a gating strategy was used to analyze the DCs, Tregs, and Bregs subsets in intestinal tissues (Figures S19A-C). As illustrated in Figure 6A and Figure S20, significant reductions in the frequency, number, and proliferation of CD103^+^CD11b^+^ DCs in the D-gut of transferrin-knockdown mice as well as Abs-treated mice were observed. Similar decreased patterns were also observed for Foxp3^+^ Tregs, RORγT^+^ Tregs (Figure 6B and Figure S21), and CD19^+^CD5^+^ Bregs (Figure 6C and Figure S22).

**Figure 6.**
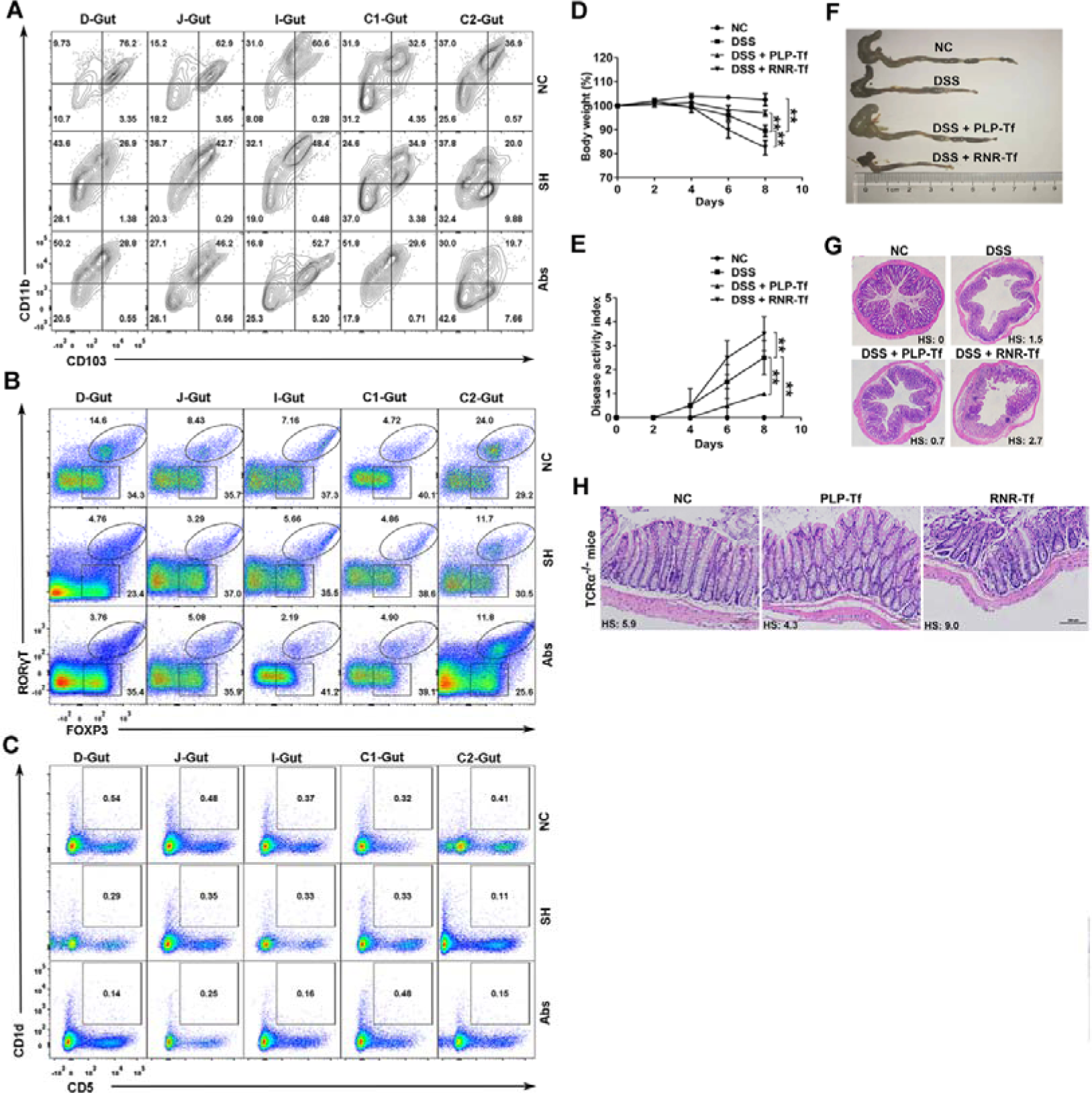
Transferrin knockdown impairs host tolerogenic responses. **(A)** DCs in different segments of gut tissue (duodenum, jejunum, ileum, caecum, and colon (D, J, I, C1, and C2)) of SPF mice (NC) and transferrin knockdown SPF mice (SH) were characterized as CD45.2^+^MHCII^+^CD11c^+^ (Figure S19A) and further subdivided into CD103^+^ DCs, CD103^+^CD11b^+^ DCs [double positive (DP) DCs], and CD11b^+^ DCs. Tregs in gut tissue **(B)** were characterized as CD45.2^+^CD4^+^ (Figure S20B) and further subdivided into Foxp3^+^ Tregs and Foxp3^+^ROROr^+^ Tregs. Bregs in gut tissue **(C)** were characterized as CD19^+^ (Figure S20C) and further subdivided into CD5^+^CD1d^+^. Lentivirus for transferrin overexpression (PLP-Tf) and retrovirus for transferrin knockdown (RNR-Tf) were injected into C57BL/6 or TCRαKO mice (16 weeks old) through the tail vein to induce transferrin overexpression and knockdown. DSS (5%) was added to drinking water of mice to induce acute colitis. Changes in body weight (%) **(D)**, disease activity index **(E)**, colon length **(F)**, and colon histopathological injury **(G)** are shown. Colon histopathological injury **(H)** at 20 weeks was also tested in TCRα KO mice with colitis. Data represent means ± SD (n = 10), \**p* < 0.05, \*\**p* < 0.01 by one-way ANOVA with Dunnett’s *post-hoc* test. Tf: transferrin. HS: histological score.

### Protective effects of transferrin on intestinal immune imbalance induced by gut microbial dysbiosis

Two experimental colitis models, which are well-known to result in intestinal dysbiosis and immune imbalance, i.e., dextran sodium sulfate (DSS) and T cell receptor α chain-deficient (TCRαKO) colitis murine models(Munyaka et al., 2016; Tamboli et al., 2004), were used to further investigate the effects of transferrin on chronic dysregulated immune response in the intestinal tract. In the DSS colitis model, weight loss in the transferrin overexpression group was lower than that in the control and transferrin knockdown group (Figure 6D). Compared with the control and transferrin knockdown mice, those with transferrin overexpression demonstrated a lower score of disease activity index (Figure 6E), longer colon length (Figure 6F and Figure S23), and alleviation of inflammation-associated histological changes (Figure 6G and Figure S24). In the TCRαKO colitis model, alleviation of inflammation-associated histological changes (Figure 6H and Figure S25) was observed in the transferrin overexpression mice compared with the control and transferrin knockdown mice.

## Methods

### Animals and ethics statement

All animal experiments were approved by the Animal Care and Use Committee of the Kunming Institute of Zoology (SMKX-2016013) and conformed to the US National Institutes of Health’s Guide for the Care and Use of Laboratory Animals (National Academies Press, 8th Edition, 2011). Germ-free (GF) and specific-pathogen-free (SPF) C57BL/6J mice (male, 8 weeks old, 10 back crosses) were purchased from the Institute of Laboratory Animal Sciences, Chinese Academy of Medical Sciences. The GF mice were maintained in sterile isolators (RSGLB-3, Huanyuzhongke, China) with autoclaved food and water until the beginning of testing. The SPF mice and some GF mice were maintained in standard SPF housing conditions for four weeks. T cell receptor α (TCRα) knockout (KO) mice (#002116, male, 16 weeks old, 12 back crosses) were purchased from the Jackson Laboratory. All mice were housed under a 12 h light-12 h dark cycle at 24 °C and tested at 10 weeks of age.

Broad-spectrum antibiotic (Abs)-treated mice (SPF C57BL/6J) were fed water with ampicillin (1 g/L), streptomycin (1 g/L), metronidazol (0.5 g/ml), and vancomycin (1 g/L) for three weeks. Lentivirus for transferrin overexpression (10^7^ transducing units (TU)), retrovirus for transferrin knockdown (10^7^ TU), or their blank viruses (10^7^ TU) were injected into SPF C57BL/6J mice through the tail vein to induce transferrin overexpression or knockdown, and the transferrin concentration was detected periodically until it was successfully overexpressed or knocked down.

### Label-free quantification

Mouse blood was obtained by mixing trisodium citrate (0.13 M) with blood at a 1:9 volume, with plasma then obtained by centrifugation at 3□000 rpm for 20 min at 4 °C. The plasma was diluted (1:2) with phosphate-buffered saline (PBS, 8 mM Na_2_HPO_4_, 0.136 M NaCl, 2 mM KH_2_PO_4_, 2.6 mM KCl, pH 7.4), followed by the addition of one volume of 2 × SDT lysate (2% SDS, 0.1M DTT, 0.1M Tris/HCL, pH 7.6). The disulfide bond of proteins was opened by vortex homogenization and ultrasonication at 100 °C for 10 min. The supernatant was collected after centrifugation at 12□000 rpm for 10 min. The acquired supernatant (100 μg, 45 μl) was alkylated with 20 mM iodoacetamide dissolved in the same buffer for 30 min at room temperature in the dark. The alkylation reaction was ended by adding additional DTT. Finally, the supernatant was treated with 1% trypsin at 37 °C overnight, then collected for mass spectrometry analysis. The data generated by the Q Exactive™ Hybrid Quadrupole-Orbitrap™ Mass Spectrometer (ThermoFisher Scientific, USA) were retrieved by Protein Discover (v2.2) using the Percolator database retrieval algorithm. The mouse proteome reference database (UniProt_Mouse_20180428.fasta) in UniProt was searched with the following parameters: Scan Event: Msaa Analyzer (FTMS), MS Order (MS2), activation type (HCD), scan typr (full); sequest HT: Enzyme (Trypsin full), Dynamic Modification (Oxidation, Acetyl, Carbamidomethyl). The search results were screened based on PSM’s Maximum Delta Cn and Maximum Rank ≥0.05. Entries retrieved from the anti-pool and contaminated proteins were deleted, and the remaining identification information was used for subsequent analyses.

### Quantitative real-time polymerase chain reaction (qRT-PCR)

Cells were harvested. RNA extraction and cDNA reverse transcription were performed using an RNA extraction kit (DP419, Tiangen, China) and reverse transcription kit (A5000, Promega, USA), respectively, as per the manufacturers’ protocols. Transferrin expression was quantified by qRT-PCR (forward primer (5’-3’): GGACGCCATGACTTTGGATG; reverse primer (5’-3’): GCCATGACAGGCACTAGACC for mouse transferrin; and forward primer (5’-3’): CCCTTAACCAATACTTCGGCTAC; reverse primer (5’-3’): GCCAAGTTCTCAAATATAGCTGAG for human transferrin). PCR was performed on a CFX-96 Touch Real-Time Detection System (Bio-Rad, USA).

### Enzyme-linked immunosorbent assay (ELISA)

Transferrin and cytokines in the samples (cell supernatant or plasma) were measured using a mouse transferrin ELISA kit (ab157724, Abcam, USA), mouse TNF-α ELISA kit (DKW12-2720-096, Dakewe Biotech, China), mouse IL-6 ELISA kit (DKW12-2060-096, Dakewe Biotech, China), mouse IL-1β ELISA kit (DKW12-2012-096, Dakewe Biotech, China), mouse IFN-β ELISA kit (SEA222Mu-96T, USCN, China), human transferrin ELISA kit (EK12012, MultiSciences, China), human TNF-α ELISA kit (DKW12-1720-096, Dakewe Biotech, China), human IL-6 ELISA kit (DKW12-1-60-096, Dakewe Biotech, China), human IFN-β ELISA kit (SEA222Hu-96T, USCN, China), and human TGF-β1 ELISA kit (DKW12-1710-096, Dakewe Biotech, China) according to the manufacturers’ instructions.

### Western blotting

Total proteins were extracted by RIPA Buffer (R0278, Sigma-Aldrich, USA) containing protease inhibitors (HY-K0010, MedChem Express, USA) and phosphatase inhibitors (HY-K0022, MedChem Express, USA). Samples were separated by 12% sodium dodecyl sulfate-polyacrylamide gel electrophoresis (SDS-PAGE) and further transferred to polyvinylidene difluoride (PVDF) membranes. The membranes were blocked with 5% BSA dissolved in TBST buffer (2.42 g/L Tris base, 8 g/L NaCl, 0.1% Tween-20 (v/v), pH 7.6) for 2 h at room temperature. After washing three times with TBST buffer, the PVDF membranes were incubated overnight in primary antibody at 4 °C, followed by thrice washing in TBST buffer and further incubation in secondary antibody at room temperature for another 1 h. Subsequently, the membranes were washed again with TBST buffer and then developed with an enhanced chemiluminescence kit (PA112, Tiangen, China) using an ImageQuant LAS 4000 mini (GE Healthcare, USA). The primary antibodies used included anti-phospho-transforming growth factor-β (TGF-β)-activating kinase 1 (TAK1) (1:1□000 dilution, AF4379-100ul, Affinity Biosciences, USA), anti-total-TAK1 (1:1□000 dilution, AF4679-50ul, Affinity Biosciences, USA), anti-phospho-inhibitory subunit of nuclear factor κB (NF-κB) (IκB) kinase α (IKKα) (1:1□000 dilution, C84E11, Cell Signaling Technology, USA), anti-total-IKKα (1:1□000 dilution, ab32041-40ul, Abcam, USA), anti-phospho-IκBα (1:1□000 dilution, AF2002-100, Affinity Biosciences, USA), anti-total-IκBα (1:2□000 dilution, 4814T, Cell Signaling Technology, USA), anti-phospho-NF-κB p65 (1:1□000 dilution, AF2006-100, Affinity Biosciences, USA), anti-total-NF-κB p65 (1:1□000 dilution, 6956T, Cell Signaling Technology, USA), anti-phospho-TRAF-associated NF-κB activator (TANK)-binding kinase 1 (TBK1) (1:1□000 dilution, AF8190-50ul, Affinity Biosciences, USA), anti-total-TBK1 (1:1□000 dilution, DF7026-50, Affinity Biosciences, USA), anti-phospho-interferon regulatory factor 3 (IRF3) (1:1□000 dilution, 29047S, Cell Signaling Technology, USA), anti-total-IRF3 (1:1□000 dilution, ab50772, Abcam, USA), anti-phospho-c-JUN N-terminal kinase (JNK) (1:1□000 dilution, 9255S, Cell Signaling Technology, USA), anti-phospho-p38 (1:1□000 dilution, 4511S, Cell Signaling Technology, USA), anti-ALDH1A2 (1:1□000, DF4422, Affinity Biosciences, USA), anti-CCL22 (1:1□000, DF7781, Affinity Biosciences, USA), anti-TGF-β1 (1:1□000, 50698-T48, Sino Biological, China), anti-IL-10 (1:1□000, GTX130513, GeneTex, USA), anti-toll-like receptor 4 (TLR4) (1:1□000, AF7017, Affinity Biosciences, China), anti-transferrin (1:5□000, ab82411, Abcam, USA), anti-GAPDH (1:3□000, T0004-50, Affinity Biosciences, USA) and anti-β-actin (1:3□000, T0022-50, Affinity Biosciences, USA).

### Bacterial and mouse DNA preparation

Bacterial isolates of gram-negative (*Escherichia coli* (ATCC 9637)) and gram-positive bacteria (*Staphylococcus aureus* (ATCC 25923) and *Listeria monocytogenes* (ATCC 19115)) were used for DNA extraction. Briefly, *E. coli* and *S. aureus* were cultured in Luria-Bertani (LB) medium (CAI-LBP03-500GM, Caisson Labs, USA). *Listeria monocytogenes* was cultured in brain heart infusion (BHI) broth (Diagnostics Pasteur, Marnes la Coquette, France). Bacterial cultures at the logarithmic phase were collected, and total bacterial DNA was extracted using a bacterial DNA extraction kit (DP302-02, Tiangen, China) as per the manufacturer’s protocols. Mouse DNA was extracted from the liver using an extraction kit (DP341-01, Tiangen, China) as per the manufacturer’s instructions.

### Cell culture

The human monocytic cell line (THP-1) and human umbilical vein endothelial cells (HUVECs) were obtained from the Conservation Genetics CAS Kunming Cell Bank (Kunming, China) and maintained in RPMI 1640 medium (Gibco Laboratories, USA) containing 10% fetal bovine serum (FBS; MRC, Australia) and M200 medium (Cascade Biologics, USA) containing 10% FBS, 2% low serum growth supplement (LSGS), soluble cluster of differentiation 14 (sCD14, 1 μg/ml), and lipopolysaccharide-binding protein (LBP, 1 μg/ml), respectively, at 37 °C and in the presence of 5% CO_2_. The mouse normal embryonic liver cell line (BNL CL.2) was also obtained from Cell Bank and maintained in Dulbecco’s modified Eagle’s medium (DMEM, Gibco Laboratories, USA) containing 10% FBS at 37 °C and under 5% CO_2_. The medium was changed every 3 d and experiments were conducted between the third and eighth passages.

### Isolation of human peripheral blood mononuclear cells (PBMCs) and polymorphonuclear neutrophils (PMNs)

Healthy human peripheral blood treated with 1.5% EDTA-Na_2_ anticoagulant agent was collected from the Kunming Blood Center, Yunnan Province, China. The PBMCs and PMNs were isolated using Polymorphprep (AS1114683, Axis-Shield, Norway) as per the manufacturer’s protocols. In brief, 5 ml of anticoagulated peripheral blood was carefully layered over 5 ml of Polymorphprep, and the samples were then centrifuged at 500 g for 30 min in a swing-out rotor at 22 °C. After centrifugation, two leucocyte bands were visible. The top band at the plasma/Polymorphprep consisted of mononuclear cells and the lower band consisted of polymorphonuclear cells. The two bands were harvested using a pipette, and the fractions of the two bands were diluted with one volume of RPMI 1640 medium. The cell suspension was transferred to a 3-ml tube and centrifuged three times at 500 g for 30 min at 22 °C. The isolated PBMCs and PMNs were maintained in RPMI 1640 medium containing 10% FBS at 37 °C and in the presence of 5% CO_2_.

### Isolation of dendritic cells from bone marrow

Femurs and tibiae of C57BL/6J mice (male, 8 weeks old) were removed and purified from the surrounding muscle tissue. The bones were left in 70% ethanol for 5 min for disinfection and washed twice with PBS. Both ends of the bones were cut with scissors. The marrow was flushed out with complete medium (CM, RPMI 1640 supplemented with 10% FBS, 2 mM L-glutamine, 1% of nonessential amino acids, 100 U/ml penicillin, and 100 μg/ml streptomycin) using a syringe with a 0.45-mm needle. Clusters within the marrow suspension were disassociated by pipetting and filtrated through a 70-μm sterilized cell strainer. Red blood cells in the suspension were lysed with Ammonium-Chloride-Potassium (ACK) lysing buffer (R1010-100, Solarbio, China). Purified bone marrow dendritic cells (BMDCs) were seeded at a concentration of 1 × l0^6^ cells/ml in RPMI 1640 medium supplemented with 20 ng/ml GM-CSF (415-ML-050, R&D Systems, USA). After 7 d of culture, the BMDCs were harvested, counted, and used for the experiments.

### Isolation of mouse splenocytes

Spleens of C57BL/6J mice (male, 8 weeks old) were removed and immediately placed in PBS. The excised spleens were sliced into small pieces and filtrated through a 70-μm sterilized cell strainer. The collected suspension was washed twice with PBS at 1□000 rpm after lysing red blood cells with ACK lysing buffer. Splenocytes were resuspended in RPMI 1640 medium and diluted to 1 × 10^6^ cells/ml for *in vitro* culture.

### Stimulation assays

THP-1 cells, HUVECs, BNL CL.2 cells, and splenocytes were seeded into 24-well plates at 4 × 10^5^ cells/well and maintained for 12 h in the corresponding medium described above. The THP-1 cells were incubated with 100 ng/ml PMA (S1819, Beyotime, China) in medium for 24 h. Different concentrations of *E. coli* lipopolysaccharides (LPS, 0.08, 0.4, and 2 μg/ml; L2630-10MG, Sigma, USA), *S. aureus* lipoteichoic acid (LTA, 0.4, 2, and 10 μg/ml; L2525-5mg, Sigma, USA), mouse DNA (0.4, 2, and 10 μg/ml), *E. coli* DNA (0.4, 2, and 10 μg/ml), *S. aureus* DNA (0.4, 2, and 10 μg/ml), or *L. monocytogenes* DNA (0.4, 2, and 10 μg/ml) were then added and incubated with the cells for 24 h at 37 °C. The cells were also treated with LPS (2 μg/ml), LTA (10 μg/ml), *E. coli* DNA (10 μg/ml), *S. aureus* DNA (10 μg/ml), or *L. monocytogenes* DNA (10 μg/ml), as described above, after pyrrolidine dithiocarbamate (PDTC, P8765-1G, Sigma, USA) pretreatment (20 μM, 30 min). The transferrin levels in cells were tested by Western blotting using anti-transferrin antibody (1:5□000 dilution, ab82411, Abcam, USA) as described above, with β-actin used as the loading control. The cell supernatant transferrin level was also tested using the ELISA kit described above. Total RNA in cells was extracted using an RNA Extraction Kit to test the expression of transferrin by qRT-PCR according to the method described above.

The THP-1 cells, HUVECs, PBMCs, PMNs, and BMDCs were seeded into 24-well plates at 4 × 10^5^ cells/well and maintained for 4 h in the corresponding medium described above. Cells were first incubated with anti-transferrin receptor (TfR) antibody (10 μg/ml, ab1086, Abcam, USA) for 30 min. Different concentrations of apo-transferrin (0.05, 0.5, and 5 μM, T4382, Sigma, USA) or holo-transferrin (0.05, 0.5, and 5 μM, T4132, Sigma, USA) were added and incubated with cells for 10 min, with LPS (2 μg/ml) then added to stimulate cells for 8 h. Supernatant levels of TNF-α, IL-6, IFN-β, and TGF-β were measured using ELISA kits, as mentioned above. Western blot analysis was used to test the phosphorylation of the TAK1, IKKα, IκBα, NF-κB, and p65 subunits of the myD88-dependent pathway and TBK1 and IRF3 of the myD88-independent pathway in THP-1 cells and HUVECs as described above. The phosphorylation levels of JNK and p38 of the mitogen-activated protein kinase (MAPK) signaling pathway were also tested.

### Surface plasmon resonance (SPR) analysis

BIAcore 2000 (GE, USA) was used to analyze the interaction between transferrin and CD14. Transferrin was first diluted (20 μg/ml) with 200 μl of sodium acetate (10 mM, pH 5), with the transferrin solution then flowed across the activated surface by NHS (N-hydroxysuccinimide) and EDC (1-ethyl-3-[3-dimethylaminopropyl] carbodiimide hydrochloride) of the CM5 sensor chip at a flow rate of 5 μl/min to couple with a CM5 sensor chip (BR100012, GE, USA) to a 500-target response value (RU). The remaining activated sites on the CM5 sensor chip were blocked by 75 μl of ethanolamine (1 M, pH 8.5). Different concentrations of CD14 (125, 250, 500, and 1000 nM; SEA518Mu-48T, Uscn, China) in Tris buffer (20 mM, pH 7.4) were applied to analyze interactions with transferrin on the surface of the CM5 sensor chip at a flow rate of 20 μl/min. The purity of all purchased proteins was greater than 98%. The equilibrium dissociation constant (*KD*) for binding, as well as the association (*Ka*) and dissociation (*Kd*) rate constants, were determined by the BIA evaluation program (GE, USA). The LPS binding region of the CD14 N-terminus (ELDDEDFRCVCNFSEPQPDWSEAFQCVSAVEVEIHAGGLN) deduced from the human CD14 sequence (GenBank number (NP_000582.1) was characterized and synthesized by GL Biochem (Shanghai, China), then analyzed by reversed phase high-performance liquid chromatography (RP-HPLC) and mass spectrometry to confirm that purity was greater than 98%. The corresponding scrambled peptide of the CD14 N-terminus (VDLSLEEENVEPRDVESQFNCESFWACDHGGCQFAPVADI) was also designed and synthesized. Different concentrations of these peptides were also applied to analyze interaction with transferrin.

The LPS antibody (10 μg/ml, MAB526Ge22-100ul, USCN, China) was dissolved in sodium acetate (10 mM, pH 5.5) and coupled with the CM5 sensor chip, as mentioned above, with LPS (1 mg/ml) then flowed across the CM5 sensor chip to combine the binding LPS antibody. CD14 (1 μM) mixed with different concentrations of transferrin (0.05, 0.5, and 5 μM) was applied to analyze the blockage of transferrin at the interaction between LPS and CD14 at a flow rate of 20 μl/min.

### Native gel electrophoresis

Basic-native gel electrophoresis was further used to analyze the interaction between transferrin and CD14. In brief, human apo-transferrin (10 μg) was first incubated with different concentrations of human CD14 (2–8 μg) in 30 μl of Tris-HCl buffer (50 mM, pH 7.4) for 10 min, and then applied to 8% precast gel (PG00810-N, Solarbio, China) to analyze complex formation between transferrin and CD14 in running buffer (0.05 M Trizma, 0.38 M glycine, pH 8.9) at 200 V constant voltage for 1 h. Staining analysis was performed with staining solution (0.25% (v/v) Coomassie Brilliant Blue R containing 50% (v/v) methanol, 40% (v/v) dH_2_O, and 10% (v/v) acetic acid) and destaining solution (consisting of 15% (v/v) methanol, 10% (v/v) acetic acid, and 75% (v/v) dH_2_O).

### Protein-protein docking

To model the transferrin-CD14 complex, we used the known structure of transferrin and CD14 for protein docking. The crystal structure of transferrin (PDB ID: 3V83) (DOI: 10.1038/nature10823) was docked to the structure of CD14 (PDB ID: 4GLP) (DOI: 10.1002/cbic.201402620) by ZDOCK. In predicting protein-protein complexes, ZDOCK considers shape complementarity, electrostatics, and desolvation free energy (https://doi.org/10.1002/prot.10389). Protein-protein docking was guided by transferrin activity and SPR experimental data (Figure 3A), where residues that disrupted binding were forced to be included in the interface and residues that did not affect binding were forced not to be included. About 2□000 structure complexes were generated and ranked according to the ZRANK scoring function. The best ZDOCK pose between the two conformations was used as a representative of transferrin-CD14 interaction.

### Recombinant expression of transferrin, CD14, and their mutants

The prokaryotic expression vector was constructed by inserting the DNA sequence encoding mature transferrin (GenBank: AAA61140.1, 679 amino acids) or transferrin mutant (R663A and K664A) between the KpnI and XhoI sites of the pSmart-I vector. The DNA encoding CD14 (NP_000582.1, 356 amino acids) or CD14 mutant (D44A, S46A and Q50A) was inserted between the BamHI and XhoI sites of the pSmart-I vector. The vectors were transformed into *E. coli* Rosetta (DE3), which was induced by 0.8 mM isopropyl β-d-thiogalactoside (IPTG) for 6 h in a 110-rpm shaker at 28 °C. After induction, *E. coli* cells were collected by centrifugation at 12□000 rpm for 15 min at 4 °C and resuspended in binding buffer (20 mM Tris-HCl, 500 mM NaCl, 5 mM imidazole, pH 7.4). The cells were then homogenized using an Ultrasonic Cell Disruption System (XINYI-IID, XinYi, China). The supernatant was collected by centrifugation at 12□000 rpm for 2 h at 4 °C. The Ni^2+^ affinity chromatography column was equilibrated in advance with binding buffer. The collected supernatant containing fusion protein was subsequently loaded on the Ni^2+^ affinity chromatography column at a flow rate of 0.7 ml/min. The bound fusion proteins were eluted with five column volumes of elution buffer (20 mM Tris-HCl, 250 mM NaCl, 500 mM imidazole, pH 7.4). The eluted fraction was resuspended in 20 mM Tris-HCl, pH 7.8, and the salt was removed using an ultrafiltration device (UFC500324, Millipore, USA).

For the release of recombinant transferrin, CD14, and their mutants, small Ubiquitin-Like Modifier (SUMO) protease (1 unit, 12588-018, Life Technologies, USA) was added to the reaction buffer (50 mM Tris-HCl, 0.2% Igepal, 1 mM dithiothreitol (DTT), pH 8.0) and maintained at 4 °C for 16 h. The reaction buffer was loaded on the Ni^2+^ affinity chromatography column again to remove SUMO protease and fusion tags. Binding properties between transferrin or transferrin mutants and CD14 or CD14 mutants were analyzed by SPR as described above.

### Fluorescent labeling of protein

Transferrin (20 mg/ml, dissolved by carbonate-bicarbonate solution (CBS, 1.59 g/L Na_2_CO_3_, 2.93 g/L NaHCO_3_, pH 9.6)) was put in a tube in a dark room. FITC buffer (5 mg/ml, dissolved with CBS; HY-66019, MCE, USA) was added into transferrin buffer drop by drop, and then the mixture was incubated for 24 h at room temperature. Sephadex G-25 column (30 × 2 cm, GE, USA) was used to purify the FITC-labeled transferrin (FITC-Tf), and PBS was used as the elution buffer. The purified FITC-labeled transferrin was freeze-dried for further use.

### Confocal microscopy

To colocalize the transferrin and CD14 complex at the membrane surface of THP-1 cells, FITC-labeled transferrin (0.5 μM) with or without anti-transferrin antibody (10 μg/ml) or control IgG (10 μg/ml) were incubated with cells for 15 min in the corresponding medium described above. After washing with PBS, cells were fixed for 15 min with 4% paraformaldehyde in PBS, blocked for 1 h at room temperature with 1% BSA, and then incubated with the antibody against CD14 (1:200 dilution, 10073-RP0, Sino Biological, China) for 1 h at 37 °C. After washing three times with PBS to remove excess primary antibodies, cells were incubated with a Cy3-labeled anti-rabbit IgG (H+L) secondary antibody (1:200 dilution, 072-01-15-06, KPL, USA) for 1 h at 37 °C. After washing with PBS to remove excess secondary antibodies, cells were stained with Prolong Gold Antifade DAPI (P36941, Life Technologies, USA) and imaged with a confocal microscope (FluoView™ 1000, Olympus, USA).

### Generation of lentiviral or retroviral vectors and virus package for transferrin overexpression or knockdown

Transferrin over-expression or knockdown vectors were constructed and HEK 293T cells (Conservation Genetics CAS Kunming Cell Bank, China) and EcoPack™ 2–293 cells (Clontech, USA) were used to package lentiviruses and retroviruses, respectively, as presented in our previous published manuscript(Tang et al., 2020).

### Murine inflammation model induced by LPS

C57BL/6J mice (male, 8 weeks old) were used. The lentivirus for transferrin overexpression (10^7^ transducing units (TU)), retrovirus for transferrin knockdown (10^7^ TU), or their blank viruses (10^7^ TU) were injected into C57BL/6J mice through the tail vein to induce transferrin overexpression or knockdown, with the transferrin concentration detected periodically. LPS (750 μg/kg) was injected into the tail vein of all mouse groups described above to induce an inflammatory response for 2 h. In the transferrin-treated group, LPS injection was performed after mouse transferrin (purity greater than 98%, T0523, Sigma, USA) administration through the tail vein for 20 min. Plasma was acquired as described above. Plasma levels of TNF-α, IL-6, IL-1β, and IFN-β were measured using the ELISA kits described above.

### Histological evaluation

The harvested liver tissues of all mouse groups described above were fixed in 4% buffered formalin overnight and dehydrated in 40% sucrose for 2 h. To assess morphological changes, frozen slices (8 μm) were then prepared using a freezing microtome (Cm3050, Leica, Germany) and stained with hematoxylin and eosin using a commercially available kit (G1120-100, Solarbio, China) as per the manufacturer’s protocols. Additional slices were processed for immunostaining of apoptotic nuclei using an apoptosis detection kit (40307ES20, Yeasen, China) following the manufacturer’s protocols and imaged by an Olympus FluoView 1000 confocal microscope.

### Segmentation of intestines

Segmentation of intestines was carried out as described previously(Esterhazy et al., 2019). The duodenum (D), jejunum (J), ileum (I), caecal colon (C1), and ascending colon (C2) were isolated. For segmentation of the small intestine, the upper 25% was taken as the duodenum, the next 50% as the jejunum, and the last 25% as the ileum.

### Isolation of lymphocytes and APCs from gut tissues

Lymphocytes and APCs were isolated as described previously(Esterhazy et al., 2019). Intestines were cut longitudinally and washed with PBS to remove the digest. Tissues were cut into 1-cm pieces and incubated in PBS with 1 μM DTT for 10 min to remove mucus. The epithelium was removed by centrifugation at 230 rpm after incubation in 20 ml of HBSS, 2% FBS, and 30 mM EDTA for 10 min at 37 °C. After washing with PBS, tissues were finely chopped and digested in 5 ml of RPMI 1640 with 2% FBS, 200 μg/ml DNaseI (10104159001, Roche, USA), and 2 mg/ml collagenase 8 (C2139, Sigma, USA) per gut segment for 45 min at 37 °C. Digests were passed through a sieve and centrifuged. Cell pellets were resuspended in 30% Percoll (MB3011, Meilunbio, China) complemented with RPMI1640, 2% FBS, and passed through a 100-μm mesh and separated by centrifugation in a discontinuous Percoll gradient (80%/30%) at 1□000 g for 25 min at room temperature. APCs and lymphocytes were isolated from the interphase, washed, and stained for flow cytometry.

### Flow cytometry

Fluorescent-dye-conjugated antibodies were purchased from BD Biosciences (USA) (anti-CD45.2, 560693; anti-CD4, 553046; anti-CD11c, 550261), eBioscience (USA) (anti-RORγT, 25-6981-82; anti-Foxp3, 12-5773-82; anti-CD1d, 17-0011-82; anti-CD5, 12-0051-82; anti-CD103, 11-1031-81; anti-MHCII, 25-5321-82), and Cell Signaling Technology (USA) (anti-CD19, 54508; anti-CD11b, 85601s). For flow cytometry of dendritic cells (DCs) (anti-CD45.2, anti-MHCII, anti-CD11c anti-CD103, and anti-CD11b), regulatory T cells (Tregs) (anti-CD45.2, anti-CD4, anti-Foxp3, and anti-RORγT), and regulatory B cells (Bregs) (anti-CD19, anti-CD1d, and anti-CD5), the isolated cells described above were surface stained using a subset of antibodies. For the detection of Foxp3 and RORγT, cells were fixed and stained using a Foxp3-staining kit (00-5523-00, eBioscience, USA) according to the manufacturer’s instructions. Dead cells were excluded by propidium iodide (PI) staining. Staining was carried out as described previously(Mortha et al., 2014) and analyzed on a BD LSRFortessa Cell Analyzer (BD Biosciences, USA).

### Ulcerative colitis model

For dextran sulphate sodium salt (DSS) treatment, C57BL6/J mice were treated with 1.5% DSS (MP Biochemicals, Canada) in drinking water for 7 d and then received regular drinking water for another 4 d. The lentiviruses or retroviruses for transferrin overexpression or knockdown were injected two weeks before DSS treatment. The mice were monitored and scored daily for body weight, presence of diarrhea, stool consistency, and bloody stools. The histological grade of colonic inflammation, colon length, and scoring of disease activity index were also tested in accordance with previously reported standards(Sang et al., 2017). For the TCRαKO model, mice were injected with the lentiviruses or retroviruses for transferrin overexpression or knockdown at 16 weeks of age, and the severity of intestinal inflammation was evaluated by histological analysis at 20 weeks as per previously described criteria(Sugimoto et al., 2007).

### Statistical analysis

The data obtained from independent experiments are presented as means ± SD. All statistical analyses were two-tailed with 95% confidence intervals (CI). The results were analyzed using one-way ANOVA with Dunnett’s *post-hoc* test or unpaired *t*-test by Prism 6 (GraphPad Software) and SPSS (SPSS Inc, USA). Differences were considered significant at *p* < 0.05.

## Discussion

This study showed that transferrin, which is the main iron transporter in serum, was up-regulated by microbial products, such as LPS, LTA, and bacterial DNA, and acted as a negative regulator of TLR4 signaling to silence host immunity, evoke immune tolerance, and maintain host intestinal homeostasis.

We found that transferrin levels in GF and Abs-treated mice were much lower than that in normal mice and that these levels returned to normal when the GF mice were fed in the SPF environment for four weeks. In addition, the change in transferrin level was consistent with that of immune tolerance-supporting factors, such as ALDH1A2, CCL22, TGF-β, and IL-10 in the gut. These results suggest that commensal microbiota or microbial products can up-regulate transferrin expression and that transferrin plays a role in maintaining intestinal homeostasis. Our study hypothesis was confirmed because 1) several types of microbial products, including LPS, LTA, and bacterial DNA, promoted transferrin expression; 2) transferrin silenced the TLR signaling complex by directly interacting with CD14, a co-receptor of many TLRs; 3) transferrin overexpression and knockdown attenuated and aggravated, respectively, the inflammatory response induced by LPS; and 4) transferrin down-regulation impaired host tolerogenic responses by dysregulating DC, Tregs, and Bregs homeostasis in the gut. The role of transferrin as an intestinal homeostasis-maintaining factor was further confirmed based on two experimental models that are widely used to study inflammatory bowel disease (IBD) and intestinal immune imbalance induced by gut microbial dysbiosis. Specifically, the DSS model, which induces epithelial damage(Eichele and Kharbanda, 2017), and the TCRαKO colitis murine model, which develops spontaneous chronic colitis similar to human ulcerative colitis at 16–20 weeks of age(Mombaerts et al., 1993). In these models, transferrin showed protective effects by suppressing the inflammatory response to colitis.

As an essential nutrient for almost all living organisms, iron participates in many important biological processes as a cofactor or prosthetic group for essential enzymes(Beard, 2001). Most human gut microbiota are devoid of iron and thus have to “steal” it from their human hosts(Weinberg, 2009). Gut microbiota have sophisticated pathways to compete with the host for iron, whereas humans and other mammals have developed strategies to cope with the bacterial piracy of iron during infection. On the one hand, higher animal hosts directly use iron to counteract microbial infection by regulating various innate host defense mechanisms, including respiratory bursts(Collins et al., 2002), iNOS-mediated production of reactive nitrogen intermediates(Alford et al., 1991; Bartfay and Bartfay, 2000), and acquired immune responses(Cunningham-Rundles et al., 2000; de Sousa and Porto, 1998; Omara and Blakley, 1994a, b; Schaible and Kaufmann, 2004; Walker and Walker, 2000). On the other hand, hosts have evolved a series of iron-mobilizing proteins to withhold iron, such as transferrin, which transports insoluble Fe^3+^ in the bloodstream and delivers it to tissues(Donovan et al., 2000; Pietrangelo, 2015). One of common adaptations that commensal microbiota share for survival is the possession of siderophores or surface receptors that directly bind transferrin and lactoferrin to acquire iron for growth(Anderson, 2018; GrayOwen and Schryvers, 1996; Kramer et al., 2019; Morgenthau et al., 2013; Qi and Han, 2018). Given no lactoferrin expression in the intestine(Ward et al., 2003) and very low amount of lactoferrin in stool (0.003-0.102 mg/day for a healthy adult subject, which is likely from hepatic bile and pancreatic juice)(Bard et al., 2003; Masson et al., 1966), transferrin seems to be a focus in the ‘iron tug of war’ between commensal bacteria and their hosts.

Microorganisms stimulate the TLR family to initiate a range of host defense mechanisms(Takeda et al., 2003). In particular, the activation of proinflammatory responses mediated by TLRs is essential for host defense, but excessive inflammation itself is maladaptive(Divanovic et al., 2005). Regulation of TLR signaling provides one point of control for excessive inflammation. Although the different types of microbial products (e.g., LPS, LTA, and DNA) studied here are recognized by different TLRs (e.g., TLR4, TLR2, and TLR9, respectively), all showed the ability to up-regulate transferrin expression, suggesting that these bacterial products may share a common pathway for the regulation of transferrin expression. We found that NF-κB plays a key role in transferrin up-regulation induced by bacterial products. NF-κB, which can be activated by members of the TLR family that recognize conserved microbial structures (i.e., LPS, LTA, and bacterial DNA)(Barton and Medzhitov, 2003), is a pivotal transcription factor involved in the regulation of a variety of proteins, including transferrin(Harizanova et al., 2005). Here, transferrin up-regulation by bacterial products was inhibited by the NF-κB inhibitor PDTC. Further investigation indicated that apo- and holo-transferrin interacted with CD14, not only as TLR co-receptors but also as pattern-recognition receptors (PRR) to play multiple roles in microbial recognition and signaling, to block signal intracellular responses after recognition of a vast array of bacterial products, thus resulting in negative regulation of TLR signaling.

In conclusion, our findings revealed that different microbiome-derived products exerted similar immunosuppressive pathways by up-regulating and beneficially using host transferrin not only as an iron supplier but also as a negative regulator of TLR signaling. Facing variable gut microbiota-host interactions, host transferrin contributes to general gut immune homeostasis by targeting CD14, a PRR and TLR co-receptor.

## Supporting information

include Figure S1-S25

## Supplementary information

Supplementary information includes Figure S1-25.

## Acknowledgments

This work was supported by the National Science Foundation of China (31930015, 21761142002, and 331372208), Chinese Academy of Sciences (XDB31000000, KFJ-BRP-008, QYZDJ-SSW-SMC012, and SAJC201606), and Yunnan Province (2015HA023 and 2018ZF001) to R.L., as well as the Ministry of Science and Technology of China (2018YFA0801403), National Science Foundation of China (31630075, 31770835, and 81770464), Chinese Academy of Sciences (XDA12040221), and Youth Innovation Promotion Association (2017432).

## Author contributions

X.T., M.F., K.X., R.C., G.W., Z.L., Z.Z, and J.M. performed the experiments and data analyses; R.L. conceived and supervised the project; R.L., X.T., and Q.L. prepared the manuscript. All authors contributed to the discussions.

## Conflicts of interest

The authors declare that they have no conflicts of interest.

