## Supplementary material for "Transferrin-dependent crosstalk between the intestinal tract and commensal microbes contributes for immune tolerance": include Figure S1-S25

**Supplementary Figure and Figure legends**

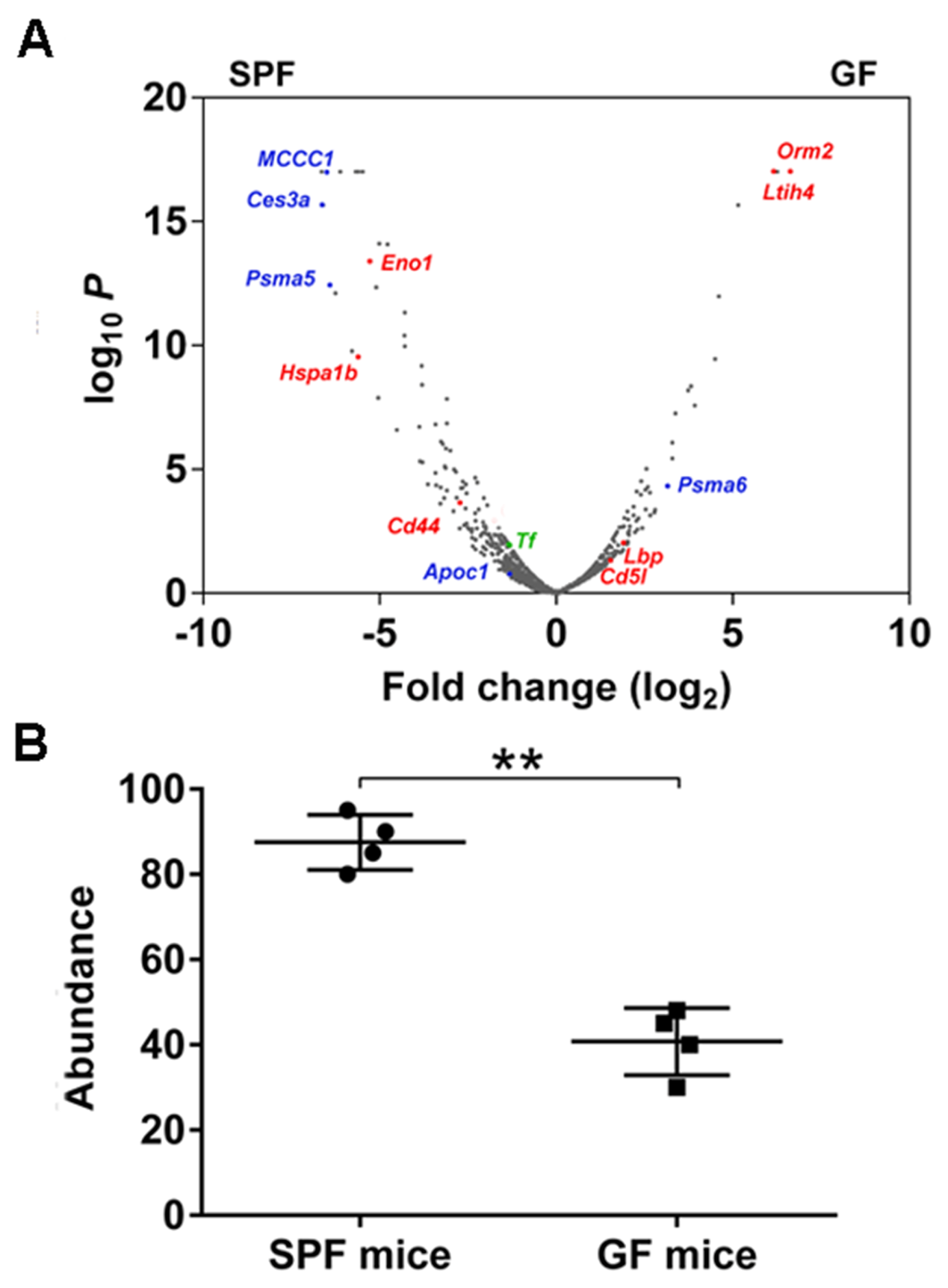

**Figure S1. Differential protein expression between specific-pathogen-free (SPF) and germ-free (GF) mice analysed by mass spectrometry-based label-free quantification. (A)** Volcano plots of all proteins quantified in SPF and GF plasma samples. Blue, metabolism-related proteins; red, immunity-related proteins; green, transferrin; All other proteins are in gray. **(B)** Jitter plot displaying relative transferrin abundance of plasma samples. Data represent means ± SD (n = 6), ***p* < 0.01 by unpaired *t*-test.

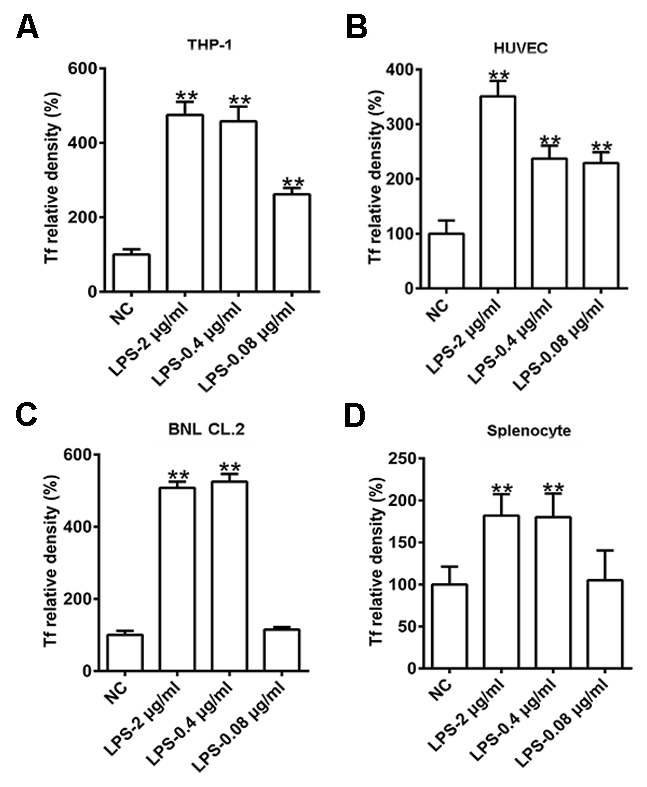

**Figure S2. Quantification analysis of transferrin up-regulated by lipopolysaccharide.** Quantifications of transferrin by Western blot analysis in ‘Figure 2A-D’ are shown. Data represent means ± SD of five independent experiments, **p* < 0.05, ***p* < 0.01 by one-way ANOVA with Dunnett’s *post-hoc* test. Tf: transferrin.

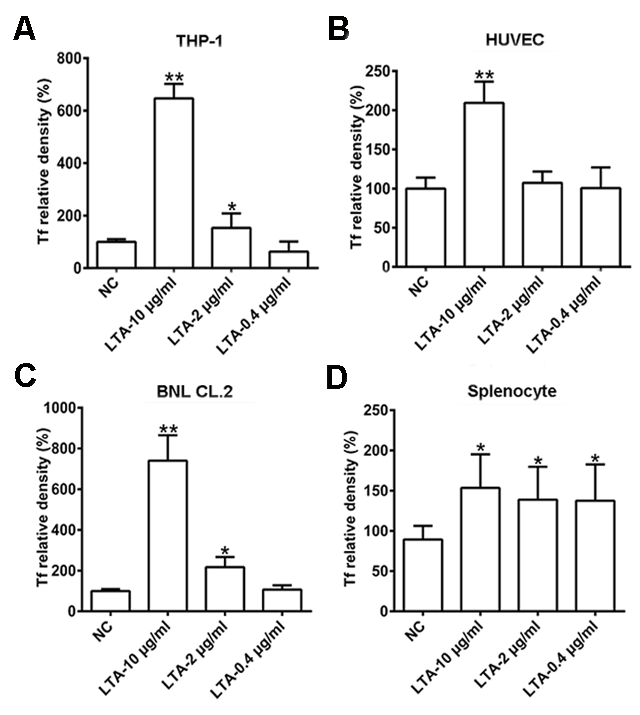

**Figure S3. Quantification analysis of transferrin up-regulated by lipoteichoic acid.** Quantifications of transferrin by Western blot analysis in ‘Figure 2E-H’ are shown. Data represent means ± SD of five independent experiments, **p* < 0.05, ***p* < 0.01 by one-way ANOVA with Dunnett’s *post-hoc* test. Tf: transferrin.

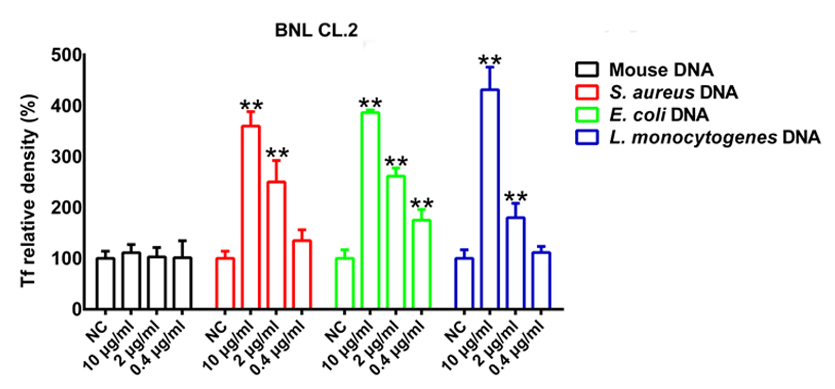

**Figure S4. Quantification analysis of transferrin up-regulated by bacterial DNA.** Quantifications of transferrin by Western blot analysis in ‘Figure 2I’ are shown. Data represent means ± SD of five independent experiments, **p* < 0.05, ***p* < 0.01 by one-way ANOVA with Dunnett’s *post-hoc* test. Tf: transferrin.

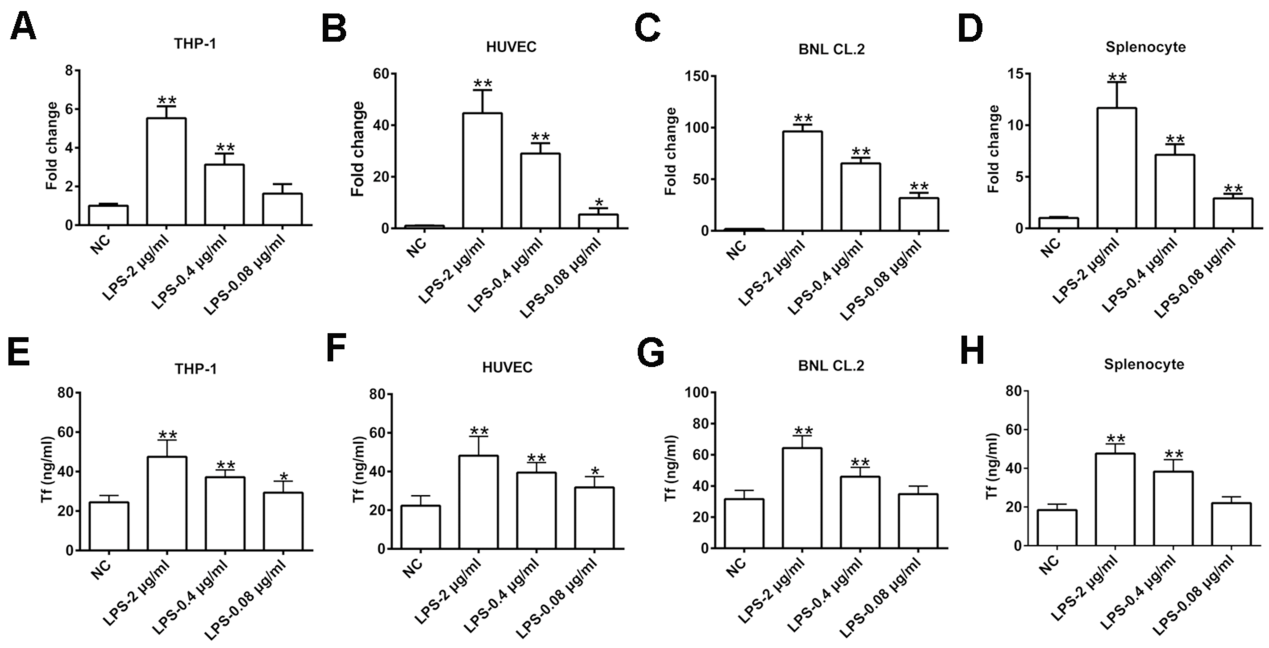

**Figure S5. Effect of lipopolysaccharide on transferrin expression.** *Transferrin* RNA expression in THP-1 cells **(A)**, HUVECs **(B)**, BNL CL.2 cells **(C)**, and splenocytes **(D)** induced by lipopolysaccharide (LPS) determined by qRT-PCR. Transferrin protein levels in THP-1 cells **(E)**, HUVECs **(F)**, BNL CL.2 cells **(G)**, and splenocytes **(H)** induced by LPS analyzed by ELISA. Data represent means ± SD of five independent experiments, **p* < 0.05, ***p* < 0.01 by one-way ANOVA with Dunnett’s *post-hoc* test.

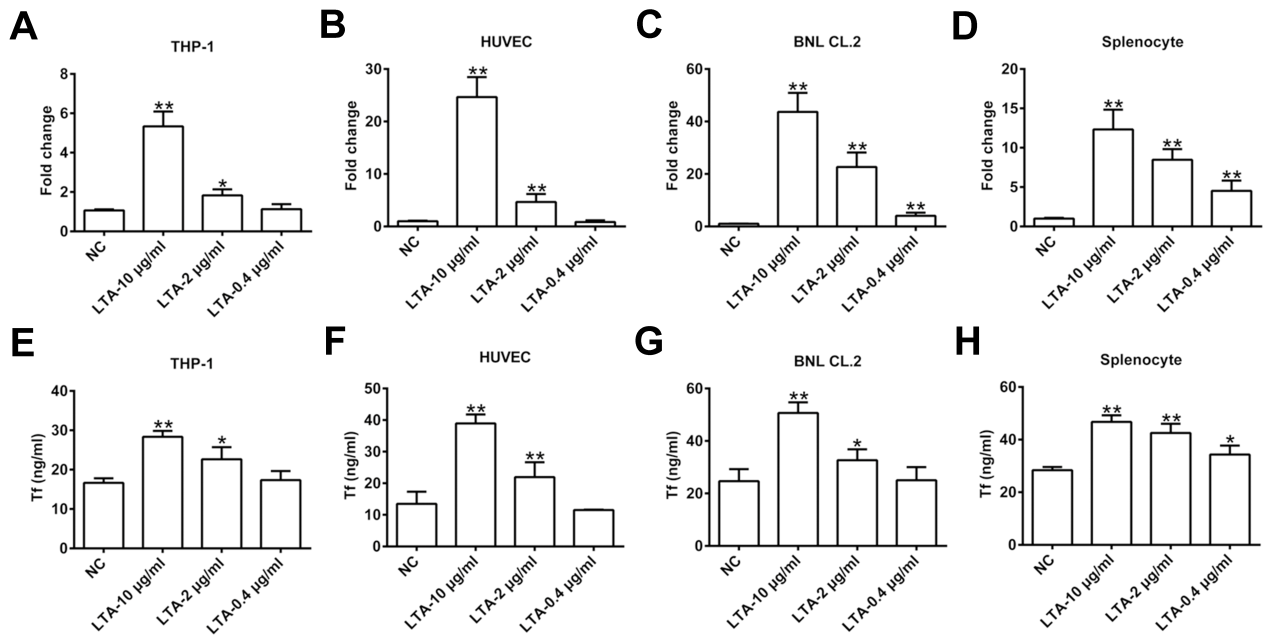

**Figure S6. Effect of lipoteichoic acid on transferrin expression.** *Transferrin* RNA expression in THP-1 cells **(A)**, HUVECs **(B)**, BNL CL.2 cells **(C)**, and splenocytes **(D)** induced by lipoteichoic acid (LTA) determined by qRT-PCR. Transferrin protein levels in THP-1 cells **(E)**, HUVECs **(F)**, BNL CL.2 cells **(G)**, and splenocytes **(H)** induced by LTA analyzed by ELISA. Data represent means ± SD of five independent experiments, **p* < 0.05, ***p* < 0.01 by one-way ANOVA with Dunnett’s *post-hoc* test.

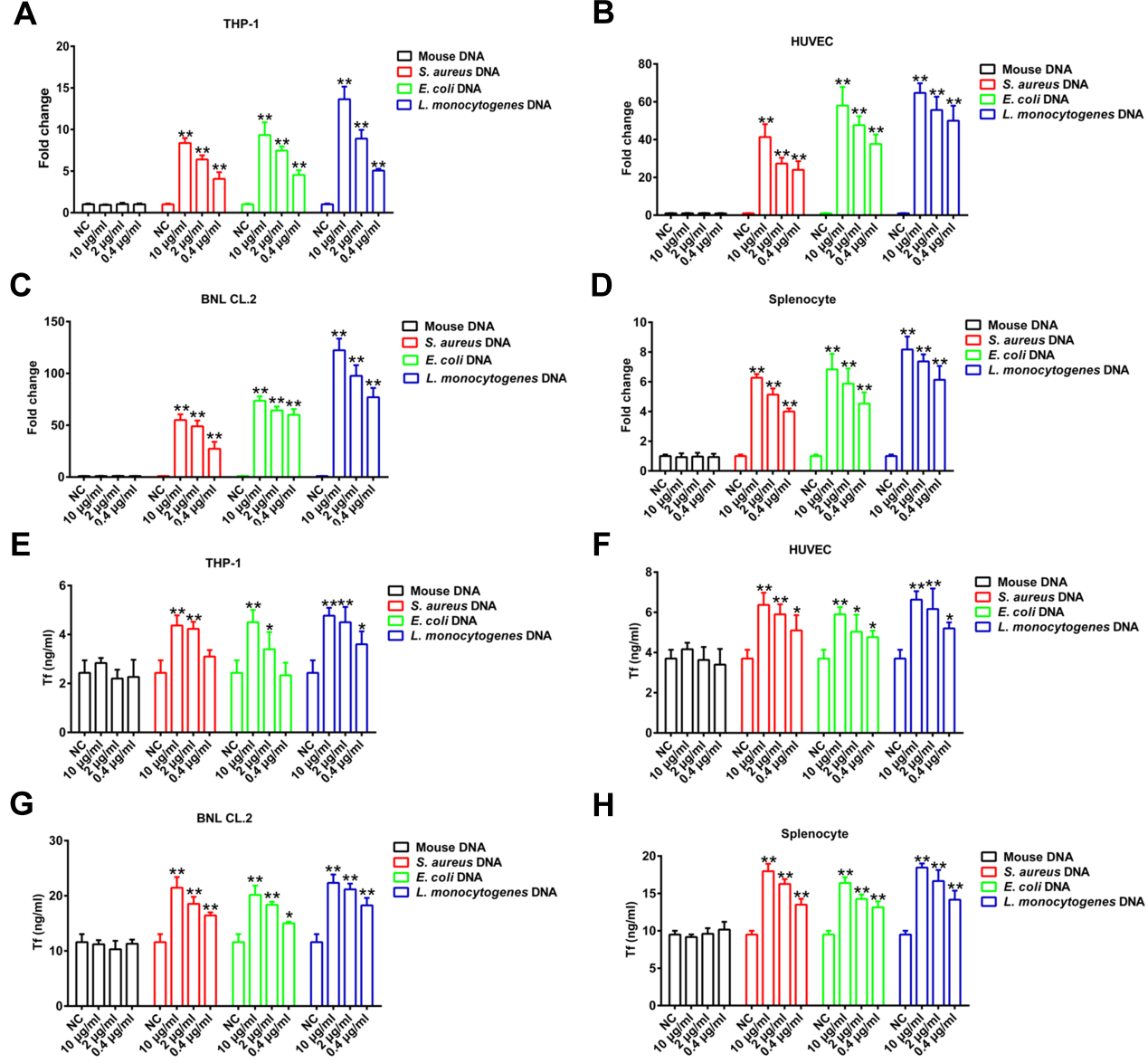

**Figure S7. Effect of bacterial DNA on transferrin expression.** *Transferrin* RNA expression in THP-1 cells **(A)**, HUVECs **(B)**, BNL CL.2 cells **(C)**, and splenocytes **(D)** induced by mouse DNA, *Escherichia coli* (*E. coli*) DNA, *Staphylococcus aureus* (*S. aureus*) DNA, and *Listeria monocytogenes* (*L. monocytogenes*) DNA determined by qRT-PCR. Transferrin protein levels in THP-1 cells **(E)**, HUVECs **(F)**, BNL CL.2 cells **(G)**, and splenocytes **(H)** induced by mouse DNA, *E. coli* DNA, *S. aureus* DNA, and *L. monocytogenes* DNA analyzed by ELISA. Data represent means ± SD of five independent experiments, **p* < 0.05, ***p* < 0.01 by one-way ANOVA with Dunnett’s *post-hoc* test.

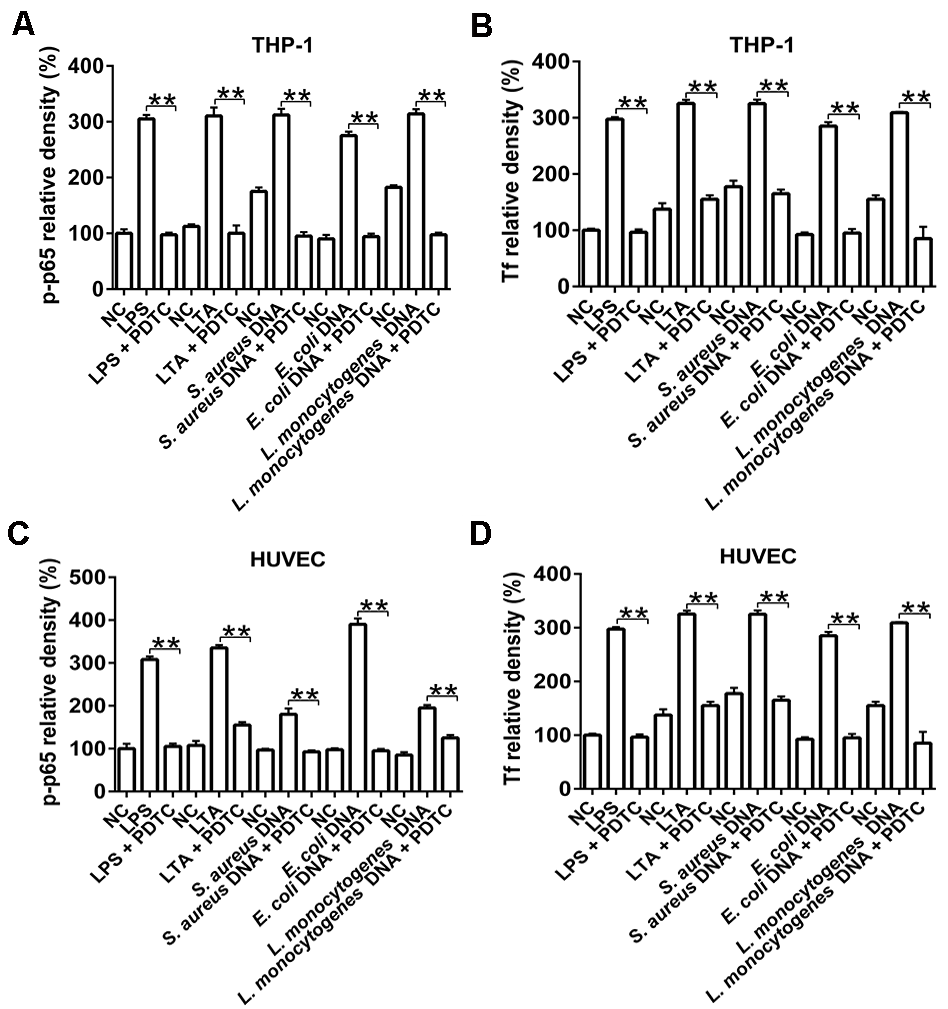

**Figure S8. Quantification analysis of inhibition of transferrin up-regulation by NF-κB inhibitor pyrrolidine dithiocarbamate.** Quantifications of inhibition of transferrin up-regulation by NF-κB inhibitor pyrrolidine dithiocarbamate by Western blot analysis in ‘Figure 2J’ are shown. Data represent means ± SD of five independent experiments, ***p* < 0.01 by unpaired *t*-test. Tf: transferrin; PDTC: pyrrolidine dithiocarbamate.

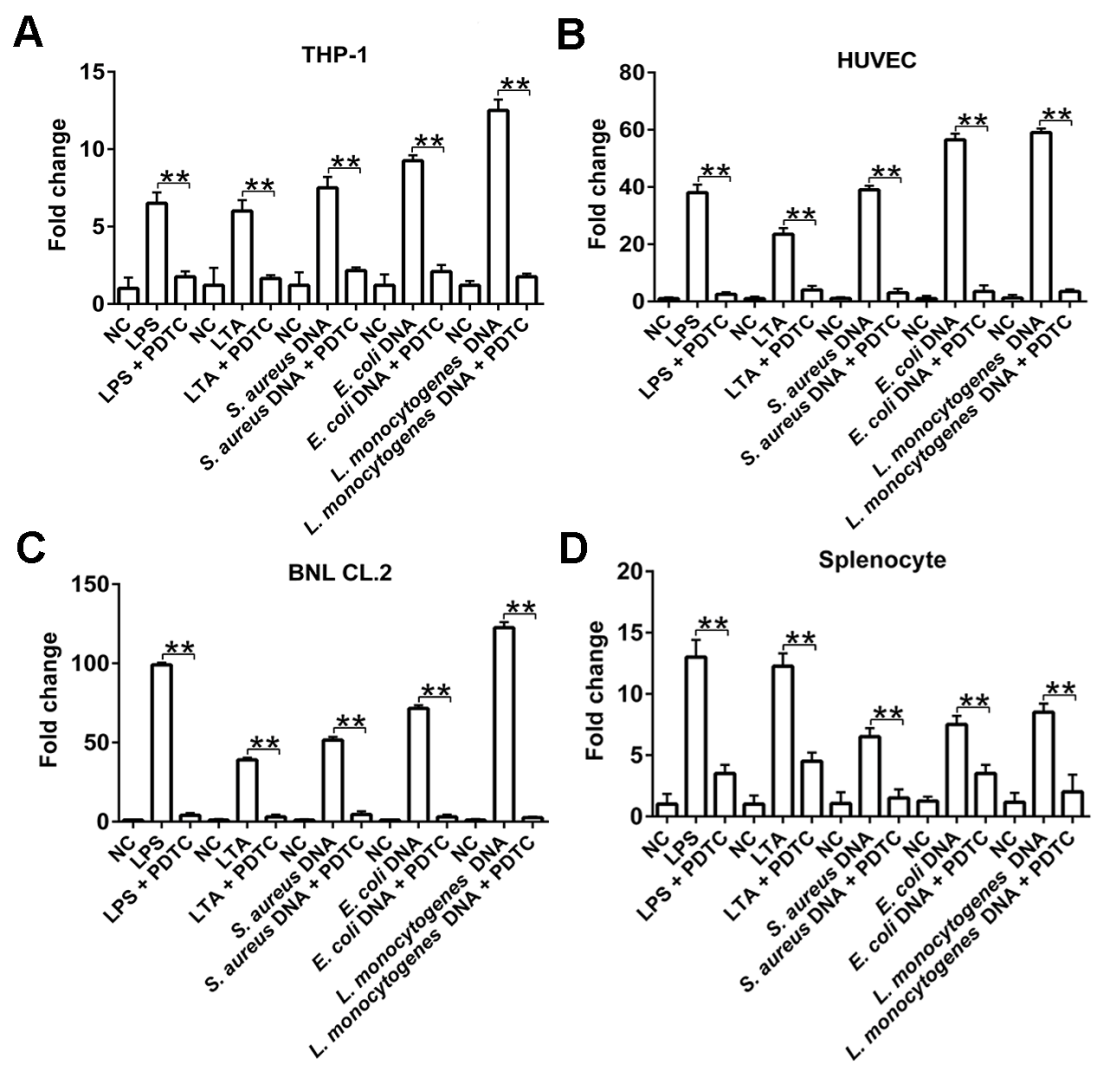

**Figure S9. Effect of pyrrolidine dithiocarbamate on transferrin expression by qRT-PCR.** Inhibition of transferrin up-regulation by NF-κB inhibitor pyrrolidine dithiocarbamate determined by qRT-PCR. Data represent means ± SD of five independent experiments, ***p* < 0.01 by unpaired *t*-test. PDTC: pyrrolidine dithiocarbamate.

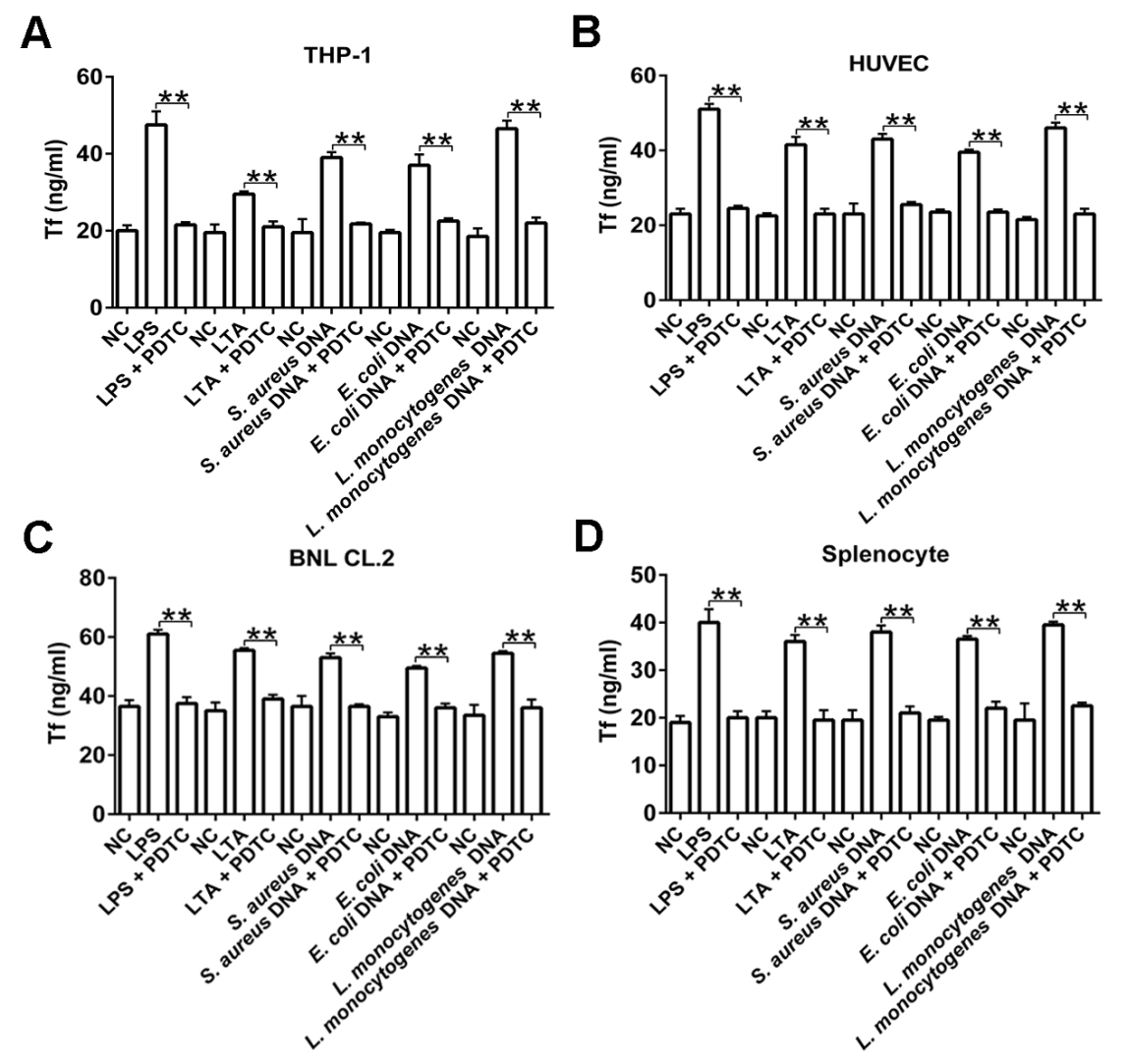

**Figure S10. Effect of pyrrolidine dithiocarbamate on transferrin expression by ELISA.** Inhibition of transferrin up-regulation by NF-κB inhibitor pyrrolidine dithiocarbamate determined by ELISA. Data represent means ± SD of five independent experiments, ***p* < 0.01 by unpaired *t*-test. Tf: transferrin; PDTC: pyrrolidine dithiocarbamate.

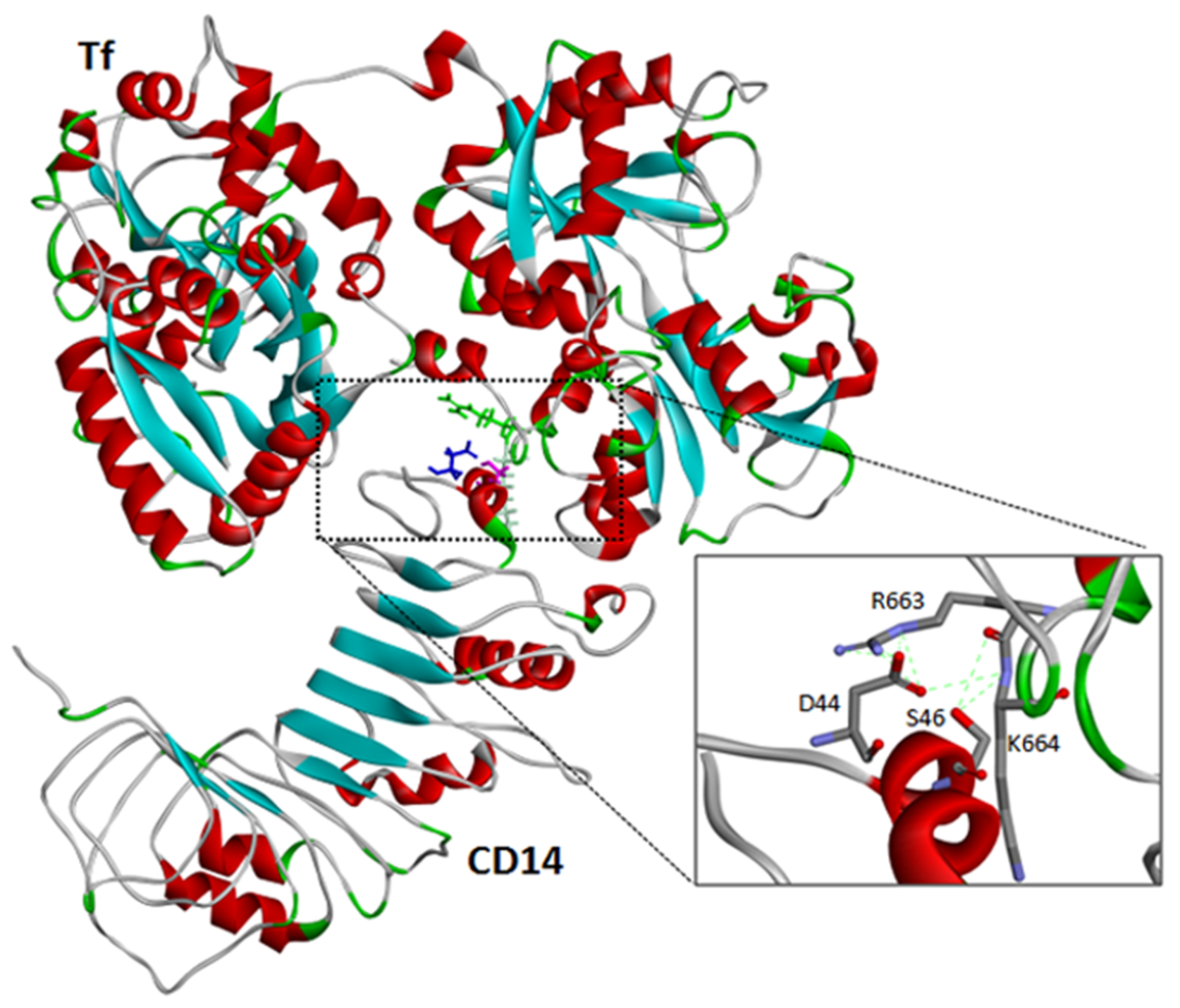

**Figure S11. Interaction of transferrin with CD14 by molecular docking.** 3D structure represents complex of transferrin and CD14. Different colors represent various secondary structure types, green: turn; red: helix; white: coil; blue: sheet. Dotted green lines in zoomed box indicate hydrogen bonds of labeled critical residues.

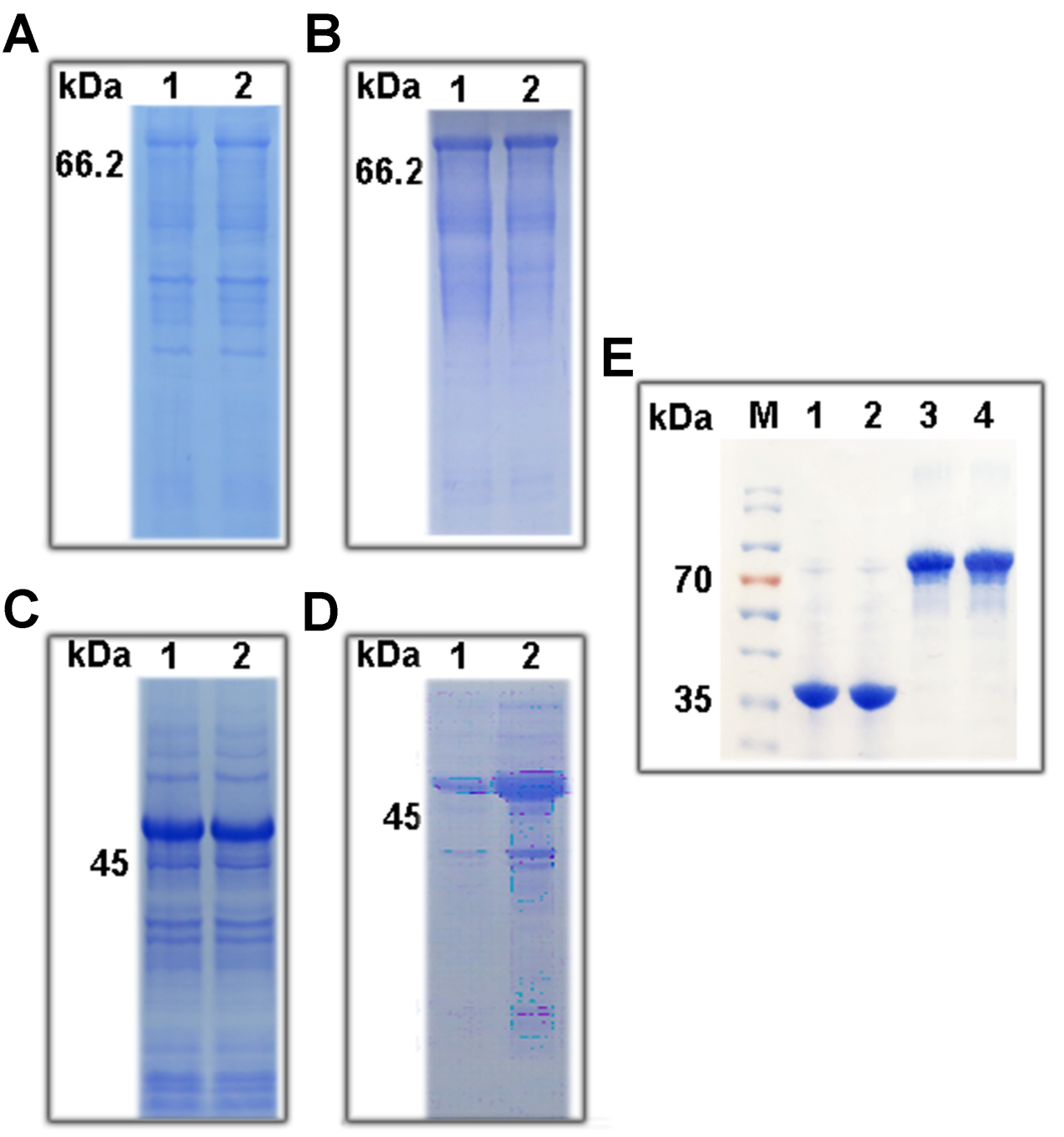

**Figure S12. Recombinant expression of transferrin, CD14, and their mutants. (A)** SDS-PAGE analysis of production of fused wild-type transferrin and its mutant after IPTG induction. Lane 1: wild-type transferrin, induced; Lane 2: transferrin mutant, induced. **(B)** Bound fusion protein of Ni^2+^ affinity chromatography column. Lane 1: fusion protein of wild-type transferrin; Lane 2: fusion protein of transferrin mutant. **(C)** SDS-PAGE analysis of production of fused wild-type CD14 and its mutant after IPTG induction. Lane 1: wild-type CD14, non-induced; Lane 2: CD14 mutant, induced. **(D)** Bound fusion protein of Ni^2+^ affinity chromatography column. Lane 1: fusion protein of wild-type CD14; Lane 2: fusion protein of CD14 mutant. **(E)** Purified transferrin, CD14, and their mutants. M: protein marker; Lane 1: purified wild-type CD14; Lane 2: purified CD14 mutant; Lane 3: purified wild-type transferrin; Lane 4: purified transferrin mutant.

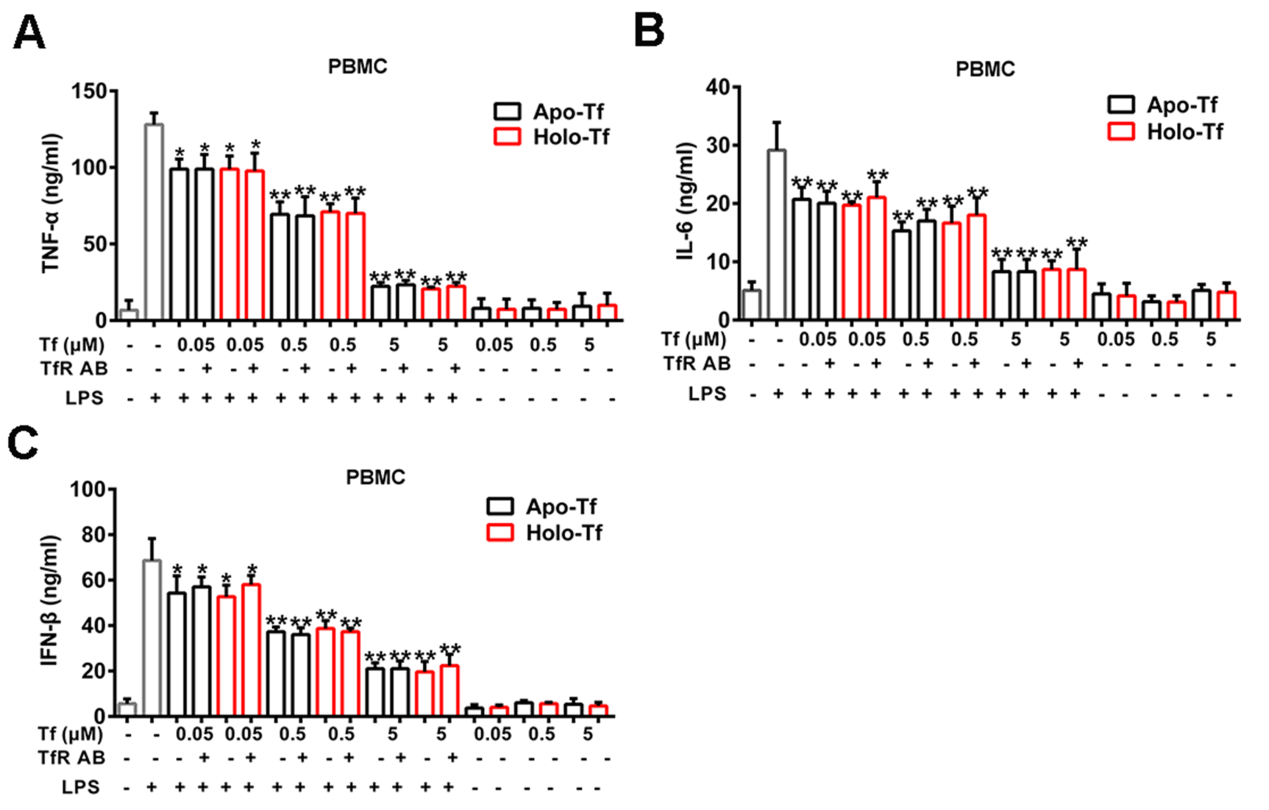

**Figure S13. Effects of apo- or holo-transferrin on production of cytokines and type I interferon in human peripheral blood mononuclear cells.** Human peripheral blood mononuclear cells (PBMCs) were stimulated in presence or absence of apo- or holo-transferrin by LPS for 8 h. Some groups of cells were first incubated with anti-transferrin receptor antibody (TfR AB, 10 μg/ml) for 30 min. Effects of apo- or holo-transferrin on TNF-α **(A)**, IL-6 **(B)**, and IFN-β **(C)** production induced by LPS in PBMCs are shown. Data represent means ± SD of five independent experiments, **p* < 0.05, ***p* < 0.01 by one-way ANOVA with Dunnett’s *post-hoc* test. Tf: transferrin.

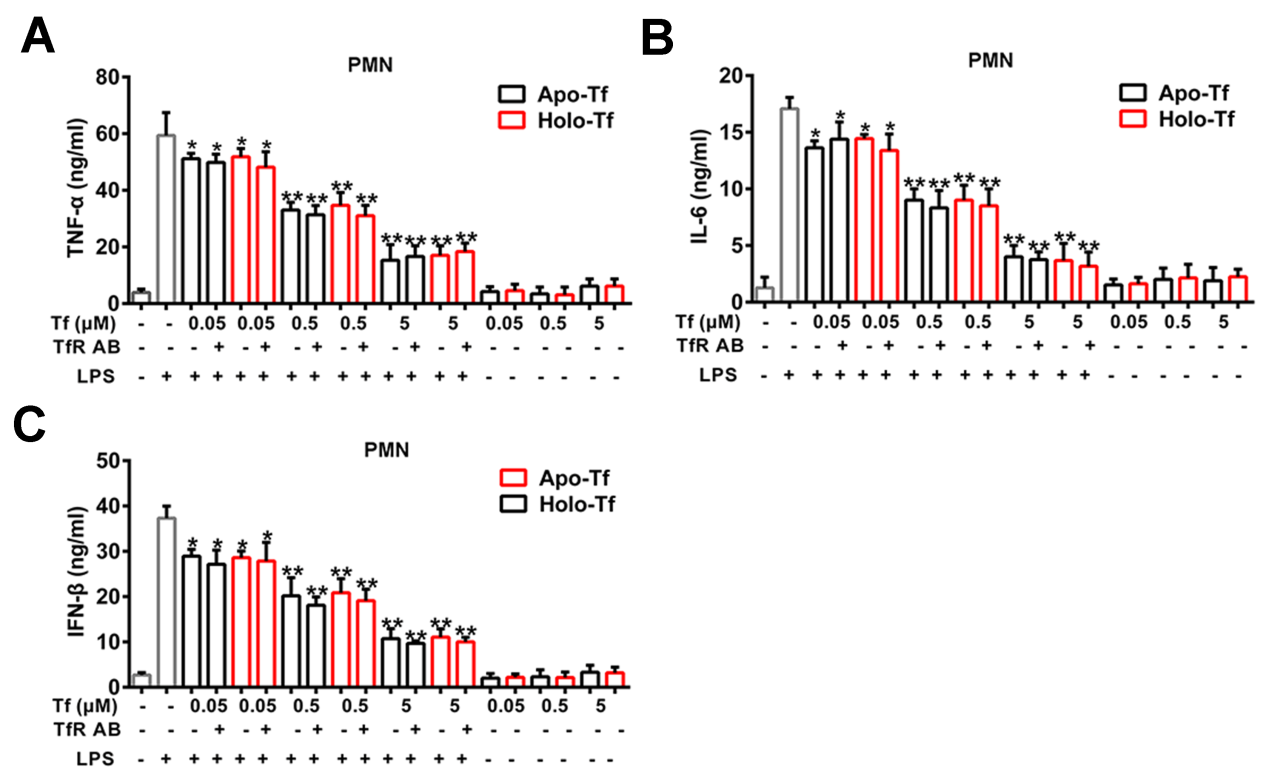

**Figure S14. Effects of apo- or holo-transferrin on production of cytokines and type I interferon in polymorphonuclear neutrophils.** Polymorphonuclear neutrophils (PMNs) were stimulated in presence or absence of apo- or holo-transferrin by LPS for 8 h. Some groups of cells were first incubated with anti-transferrin receptor antibody (TfR AB, 10 μg/ml) for 30 min. Effects of apo- or holo-transferrin on TNF-α **(A)**, IL-6 **(B)**, and IFN-β **(C)** production induced by LPS in PMNs are shown. Data represent means ± SD of five independent experiments, **p* < 0.05, ***p* < 0.01 by one-way ANOVA with Dunnett’s *post-hoc* test. Tf: transferrin.

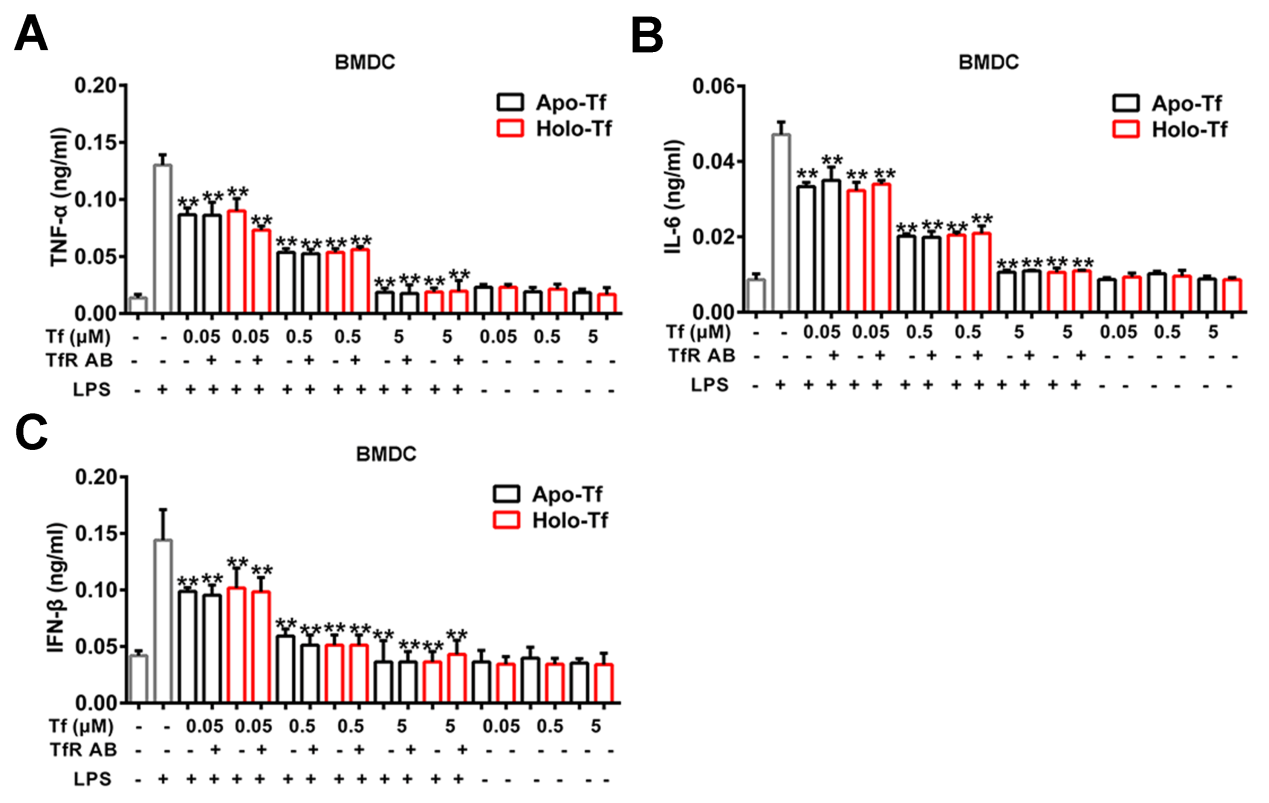

**Figure S15. Effects of apo- or holo-transferrin on production of cytokines and type I interferon in mouse bone marrow dendritic cells.** Mouse bone marrow dendritic cells (BMDCs) were stimulated in presence or absence of apo- or holo-transferrin by LPS for 8 h. Some groups of cells were first incubated with anti-transferrin receptor antibody (TfR AB, 10 μg/ml) for 30 min. Effects of apo- or holo-transferrin on TNF-α **(A)**, IL-6 **(B)**, and IFN-β **(C)** production induced by LPS in BMDC are shown. Data represent means ± SD of five independent experiments, **p* < 0.05, ***p* < 0.01 by one-way ANOVA with Dunnett’s *post-hoc* test. Tf: transferrin.

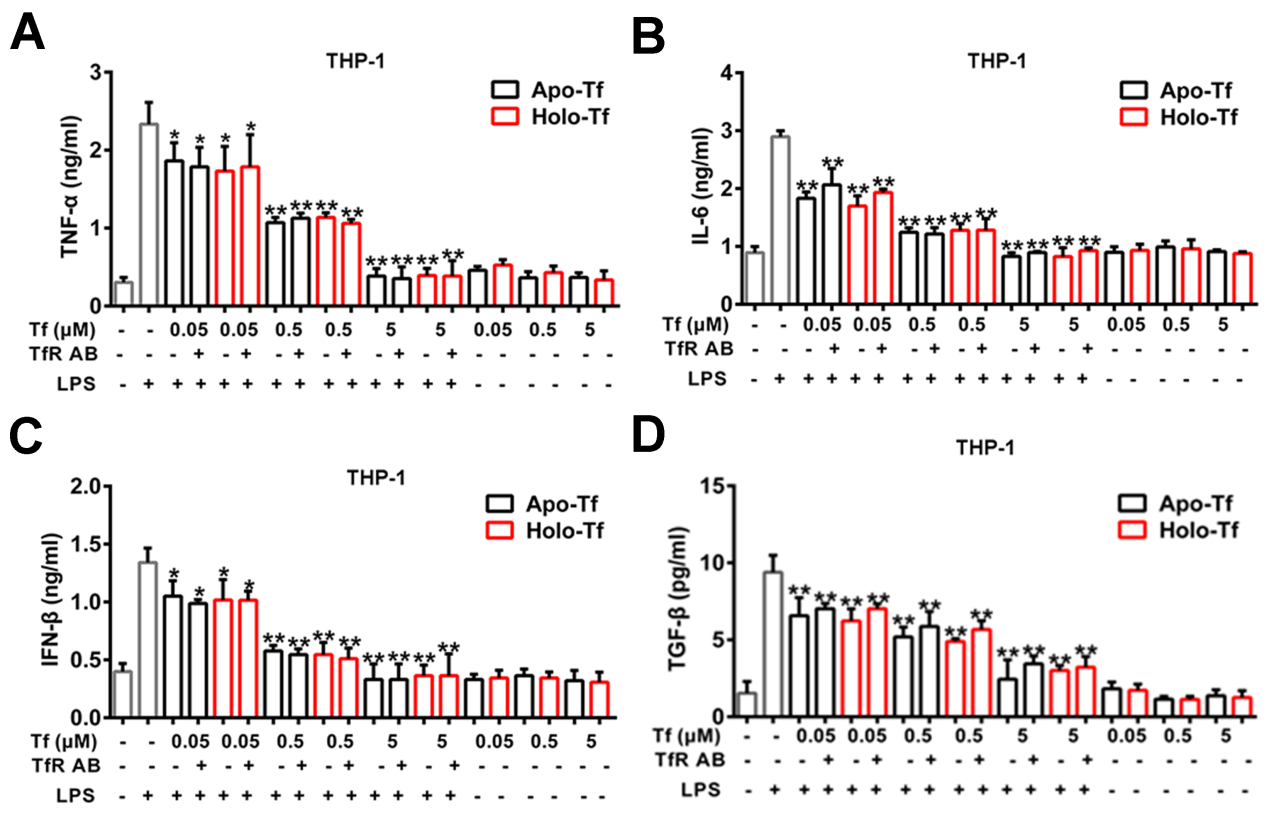

**Figure S16. Effects of apo- or holo-transferrin on production of cytokines and type I interferon in human monocytic cell line.** Human monocytic cells (THP-1) were stimulated in presence or absence of apo- or holo-transferrin by LPS for 8 h. Some groups of cells were first incubated with anti-transferrin receptor antibody (TfR AB, 10 μg/ml) for 30 min. Effects of apo- or holo-transferrin on TNF-α **(A)**, IL-6 **(B)**, IFN-β **(C)**, and TGF-β **(D)** production induced by LPS in THP-1 cells are shown. Data represent means ± SD of five independent experiments, **p* < 0.05, ***p* < 0.01 by one-way ANOVA with Dunnett’s *post-hoc* test. Tf: transferrin.

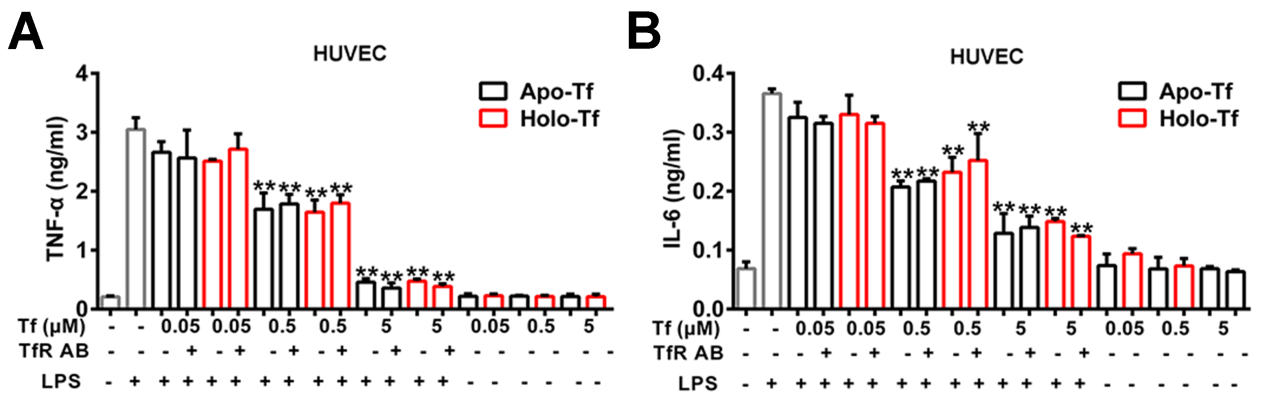

**Figure S17. Effects of apo- or holo-transferrin on production of cytokines and type I interferon in human umbilical vein endothelial cells.** Human umbilical vein endothelial cells (HUVECs) were stimulated in presence or absence of apo- or holo-transferrin by LPS for 8 h. Some groups of cells were first incubated with anti-transferrin receptor antibody (TfR AB, 10 μg/ml) for 30 min. Effects of apo- or holo-transferrin on TNF-α **(A)** and IL-6 **(B)** production induced by LPS in HUVECs are shown. Data represent means ± SD of five independent experiments, **p* < 0.05, ***p* < 0.01 by one-way ANOVA with Dunnett’s *post-hoc* test. Tf: transferrin.

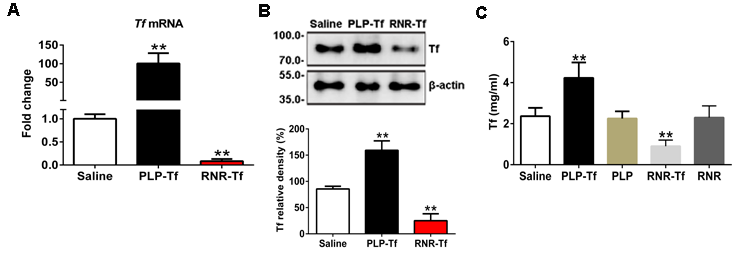

**Figure S18: Construction of transferrin overexpression or knockdown vectors.** **(A)** *Transferrin* mRNA levels of BNL CL.2 cells after transfection of overexpression or knockdown plasmid of transferrin determined by qRT-PCR. **(B)** Transferrin levels in BNL CL.2 cells determined by Western blotting (top, Lane 1: control (Saline), Lane 2: overexpression (PLP-Tf), Lane 3: knockdown (RNR-Tf)). Quantification of Western blots is also shown (bottom). Data represent means ± SD of six independent experiments, ***p* < 0.01 by one-way ANOVA with Dunnett’s *post-hoc* test. Tf: transferrin. **(C)** Plasma concentrations of transferrin in three groups of C57BL/6J mice (PLP-Tf and its blank PLP, RNR-Tf and its blank RNR, and control mice (Saline)). Data represent means ± SD (n = 10), ***p* < 0.01 by one-way ANOVA with Dunnett’s *post-hoc* test. Tf: transferrin.

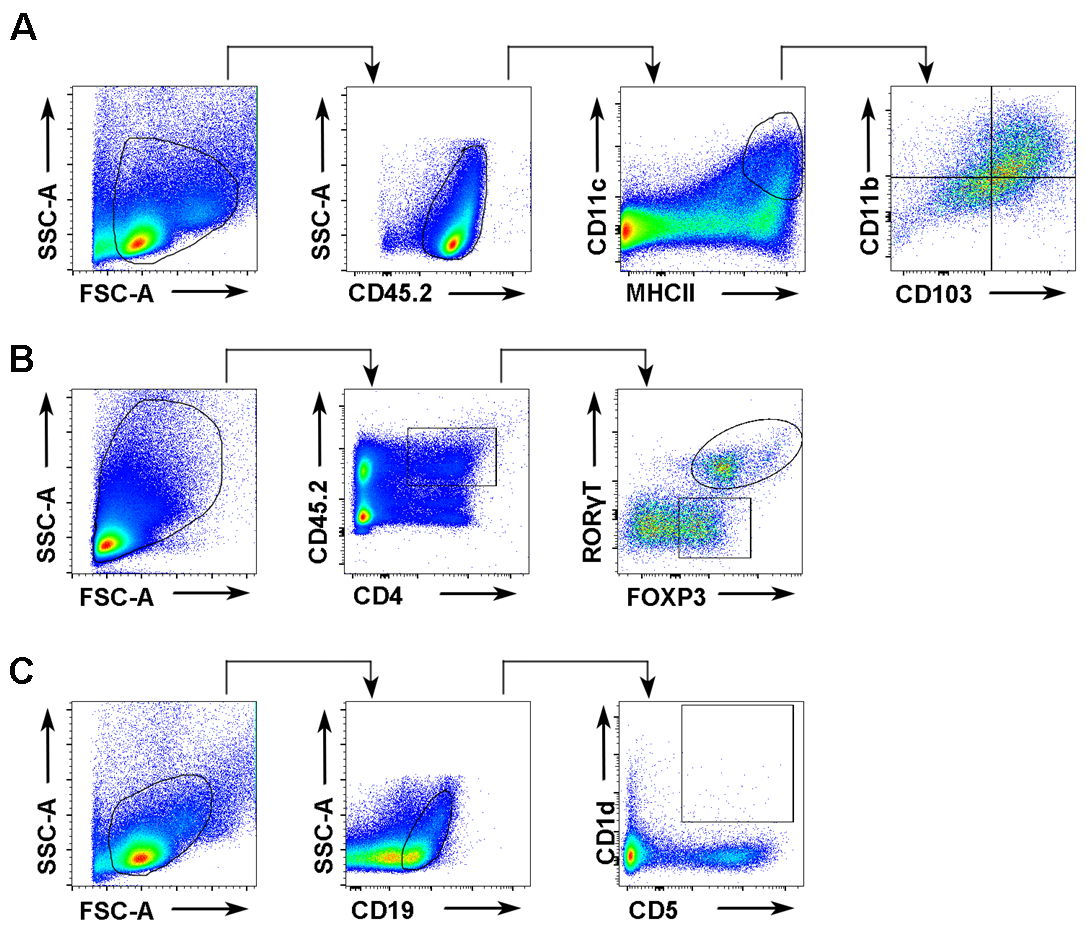

**Figure S19:** **Gating strategy to analyze intestinal tissues and lymph nodes of DCs, Tregs and Bregs subsets.** **(A)** Cells isolated from gut tissues and gut-draining lymph nodes were stained with anti-45.2, anti-MHCII, anti-CD11c, anti-CD11b, and anti-CD103 antibodies. CD103^+^ DCs were identified as CD45.2^+^MHCII^+^CD11c^+^CD11b^-^CD103^+^, double positive (DP) DCs were identified as CD45.2^+^MHCII^+^CD11c^+^CD11b^+^CD103^+^, and CD11b^+^ DCs were identified as CD45.2^+^MHCII^+^CD11c^+^CD11b^+^CD103^-^. **(B)** Cells isolated from gut tissues and gut-draining lymph nodes were stained with anti-45.2, anti-CD4, anti-Foxp3, and anti-RORγT antibodies. Foxp3^+^ Tregs were identified as CD45.2^+^CD4^+^Foxp3^+^RORγT^-^ and Foxp3^+^RORγT^+^ Tregs were identified as CD45.2^+^CD4^+^Foxp3^+^RORγT^+^. **(C)** Bregs were identified as CD19^+^CD5^+^CD1d^+^ from gut tissues and gut-draining lymph nodes.

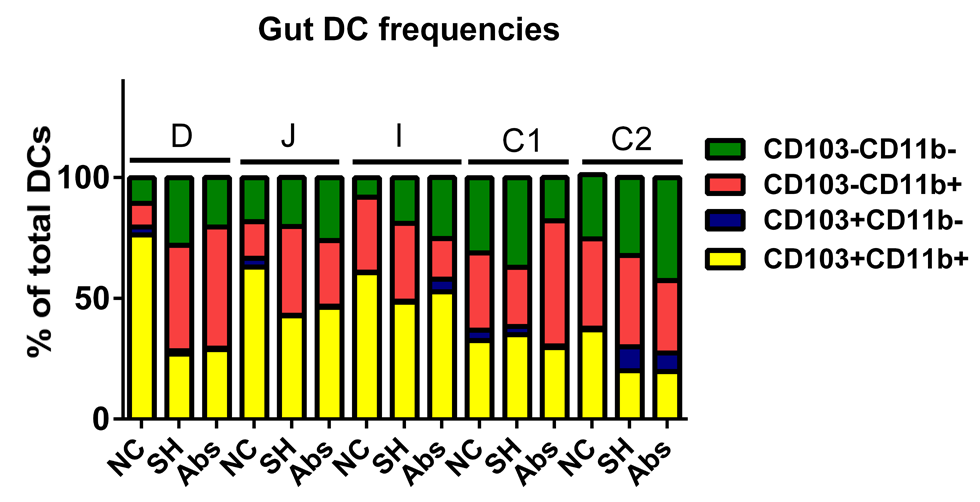

**Figure S20. Quantification analysis of DC cells.** Quantifications of CD103^+^ DCs, DP DCs, and CD11b^+^ DCs in ‘Figure 6A’ are shown.

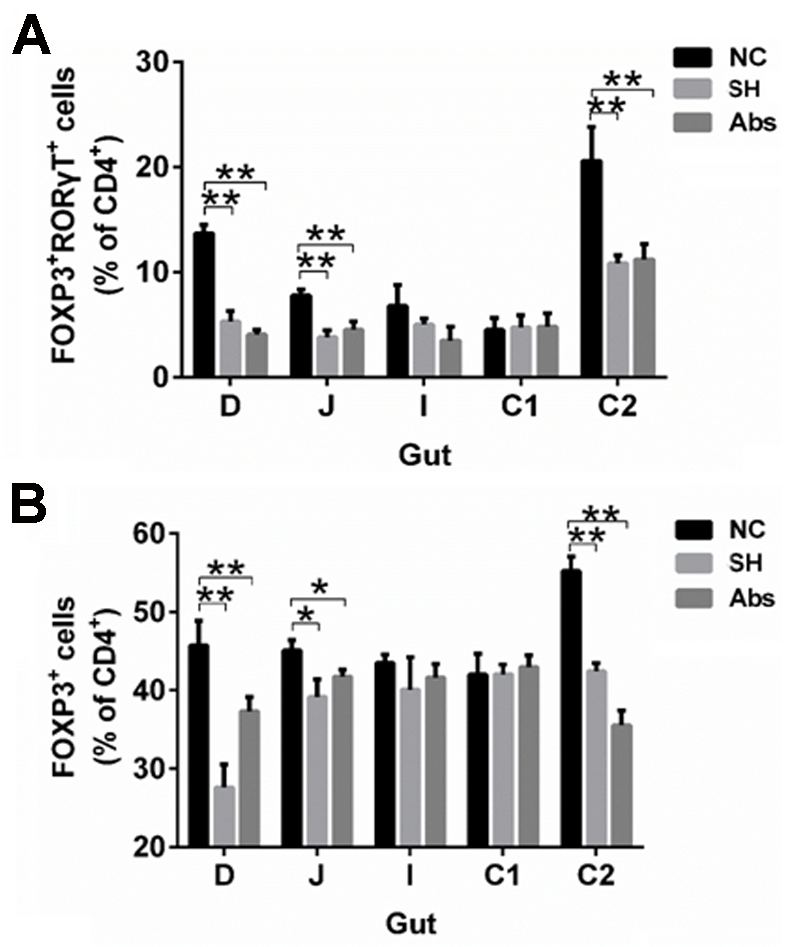

**Figure S21. Quantification analysis of Tregs.** Quantifications of Foxp3^+^ Tregs and Foxp3^+^ROROT^+^ Tregs in ‘Figure 6B’ are shown. Data represent means ± SD (n = 10), **p* < 0.05, ***p* < 0.01 by one-way ANOVA with Dunnett’s *post-hoc* test.

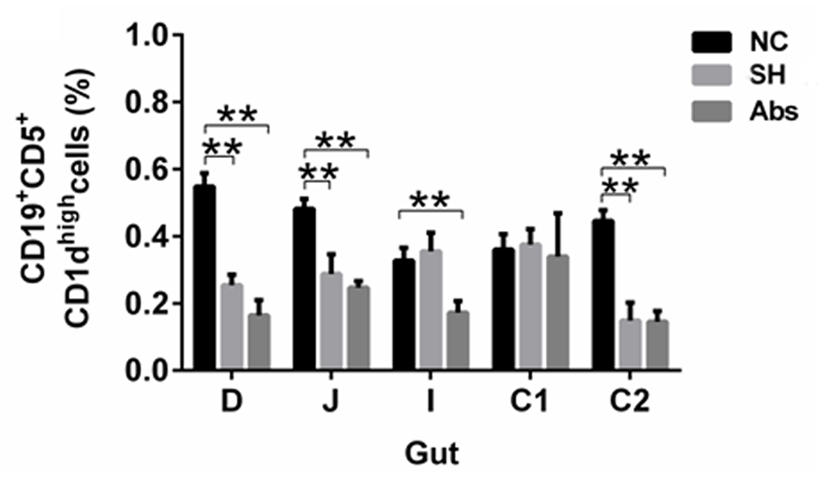

**Figure S22. Quantification analysis of Breg.** Quantifications of Bregs in ‘Figure 6C’ are shown. Data represent means ± SD (n = 10), **p* < 0.05, ***p* < 0.01 by one-way ANOVA with Dunnett’s *post-hoc* test.

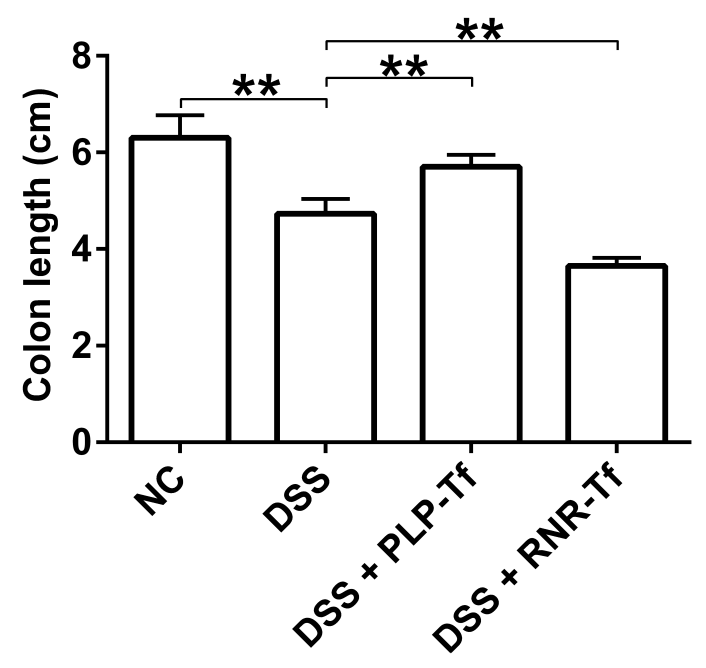

**Figure S23. Quantification analysis of colon length.** Quantification of colon length in ‘Figure 6F’ is shown. Data represent means ± SD (n = 10), **p* < 0.05, ***p* < 0.01 by one-way ANOVA with Dunnett’s *post-hoc* test.

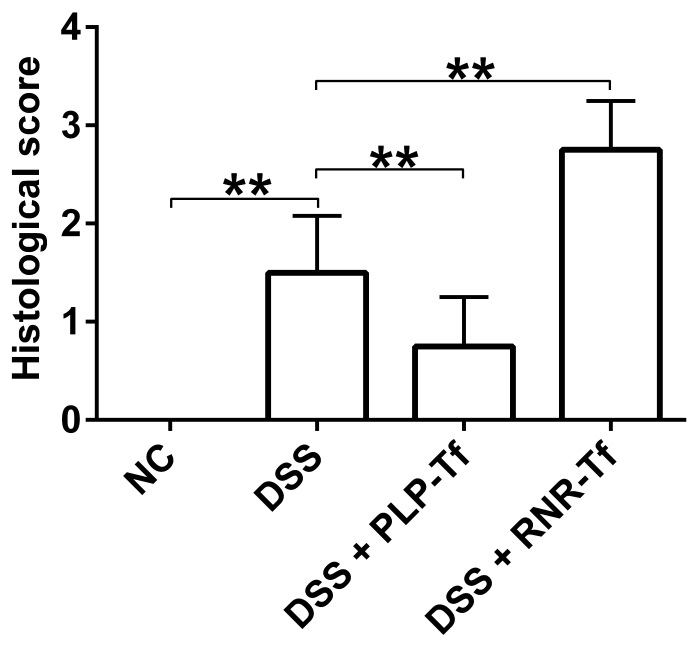

**Figure S24. Quantification analysis of hematoxylin and eosin staining.** Quantification of colon histopathological injury in ‘Figure 6G’ is shown. Data represent means ± SD (n = 10), **p* < 0.05, ***p* < 0.01 by one-way ANOVA with Dunnett’s *post-hoc* test.

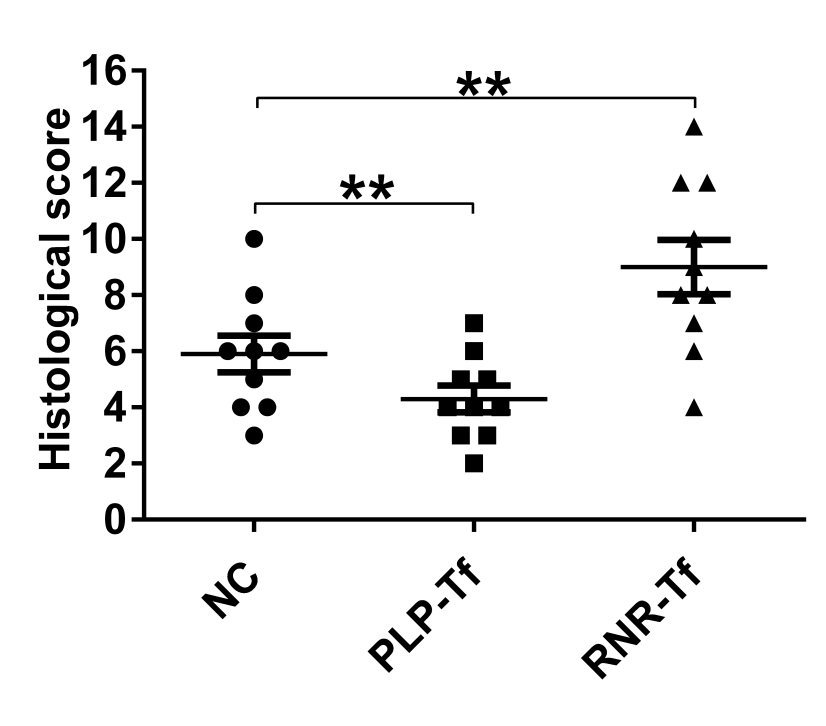

**Figure S25. Quantification analysis of hematoxylin and eosin staining.** Quantification of colon histopathological injury in ‘Figure 6H’ is shown. Data represent means ± SD (n = 10), **p* < 0.05, ***p* < 0.01 by one-way ANOVA with Dunnett’s *post-hoc* test.
